# Population structure of modern-day Italians reveals patterns of ancient and archaic ancestries in Southern Europe

**DOI:** 10.1101/494898

**Authors:** A. Raveane, S. Aneli, F. Montinaro, G. Athanasiadis, S. Barlera, G. Birolo, G. Boncoraglio, AM. Di Blasio, C. Di Gaetano, L. Pagani, S. Parolo, P. Paschou, A. Piazza, G. Stamatoyannopoulos, A. Angius, N. Brucato, F. Cucca, G. Hellenthal, A. Mulas, M. Peyret-Guzzon, M. Zoledziewska, A. Baali, C. Bycroft, M. Cherkaoui, C. Dina, JM. Dugoujon, P. Galan, J. Giemza, T. Kivisild, M. Melhaoui, M. Metspalu, S. Myers, LM. Pereira, FX. Ricaut, F. Brisighelli, I. Cardinali, V. Grugni, H. Lancioni, V.L. Pascali, A. Torroni, O. Semino, G. Matullo, A. Achilli, A. Olivieri, C. Capelli

**Author notes:** Correspondence (A.R.), (S.A.), (F.M.) and (C.C.). These authors contributed equally to this work (A. Raveane, S. Aneli, F. Montinaro). Co-senior authors (G. Matullo, A. Achilli, A. Olivieri, C. Capelli).

## Abstract

European populations display low genetic diversity as the result of long term blending of the small number of ancient founding ancestries. However it is still unclear how the combination of ancient ancestries related to early European foragers, Neolithic farmers and Bronze Age nomadic pastoralists can fully explain genetic variation across Europe. Populations in natural crossroads like the Italian peninsula are expected to recapitulate the overall continental diversity, but to date have been systematically understudied. Here we characterised the ancestry profiles of modern-day Italian populations using a genome-wide dataset representative of modern and ancient samples from across Italy, Europe and the rest of the world. Italian genomes captured several ancient signatures, including a non-steppe related substantial ancestry contribution ultimately from the Caucasus. Differences in ancestry composition as the result of migration and admixture generated in Italy the largest degree of population structure detected so far in the continent and shaped the amount of Neanderthal DNA present in modern-day populations.

**One sentence summary:** Ancient and historical admixture events shaped the genetic structure of modern-day Italians, the ancestry profile of Southern European populations and the continental distribution of Neanderthal legacy.

## Introduction

Our understanding of the events that shaped European genetic variation has been redefined by the availability of ancient DNA (aDNA). In particular, it has emerged that, in addition to the contributions of early hunter-gatherer populations, major genetic components can be traced back to Neolithic (*1*–*4*) and Bronze Age expansions (*3*, *5*).

The arrival of farming in Europe from Anatolia led to a partial replacement via admixture of autochthonous and geographically structured hunter-gatherers, a process that generated individuals genetically close to present-day Sardinians (*2, 4, 6, 7*). During the Bronze Age the dispersal of a population related to the pastoralist nomadic Yamnaya from the Pontic-Caspian steppe area dramatically impacted the genetic landscape of the continent, particularly of Northern and Central Europe (*3, 5, 8*). This migration, supported by archaeological and genetic data, has also been putatively linked to the spread of the Indo-European languages in Europe and the introduction of several technological innovations in peninsular Eurasia (*9*). Genetically, ancient steppe populations have been described as a combination of Eastern and Caucasus Hunter Gatherer/Iran Neolithic ancestries (EHG and CHG/IN) (*6*), whose genetic signatures in the population of Central and Northern Europe were introduced via admixture. However, the analysis of aDNA from Southern East Europe identified the existence of additional contributions ultimately from the Caucasus (*10, 11*) and suggested a more complex ancient ancestry composition for Europeans (*6*).

The geographic location of Italy, enclosed between continental Europe and the Mediterranean Sea, makes the Italian people relevant for the investigation of continent-wide demographic events, to complement and enrich the information provided by aDNA studies. In order to characterise the ancestry profile of modern-day populations and test the validity of the three-ancestries model across Europe (related to early European foragers, Neolithic farmers and Bronze Age nomadic pastoralists), we characterised the genetic variability of present-day Italians and other Europeans in terms of their ancient ancestry composition as the result of migration and admixture. In doing so, we assembled and analyzed a comprehensive genome-wide SNP dataset composed by 1,616 individuals from all the 20 Italian administrative regions and more than 140 worldwide reference populations, for a total of 5,192 modern-day samples (fig. S1, table S1), to which we added genomic data available for ancient individuals (data file S1).

## Results

### Distinctive genetic structure in Italy

We initially investigated patterns of genetic differentiation in Italy and surrounding regions by exploring the information embedded in SNP-based haplotypes of modern samples (Full Modern Dataset, FMD, including 218,725 SNPs). The phased genome-wide dataset was analysed using the CHROMOPAINTER (CP) and fineSTRUCTURE (fS) pipeline (*12*, *13*) (Supplementary materials) to generate a tree of groups of individuals with similar “copying vectors” (clusters, Fig. 1A). The fraction of pairs of individuals placed in the same cluster across multiple runs was on average 0.95 for Italian clusters and 0.96 across the whole set of clusters (see Materials and Methods, Supplementary materials). Related non-European clusters were merged into larger groups in subsequent analyses (see Materials and Methods, Supplementary materials).

**Fig. 1.**
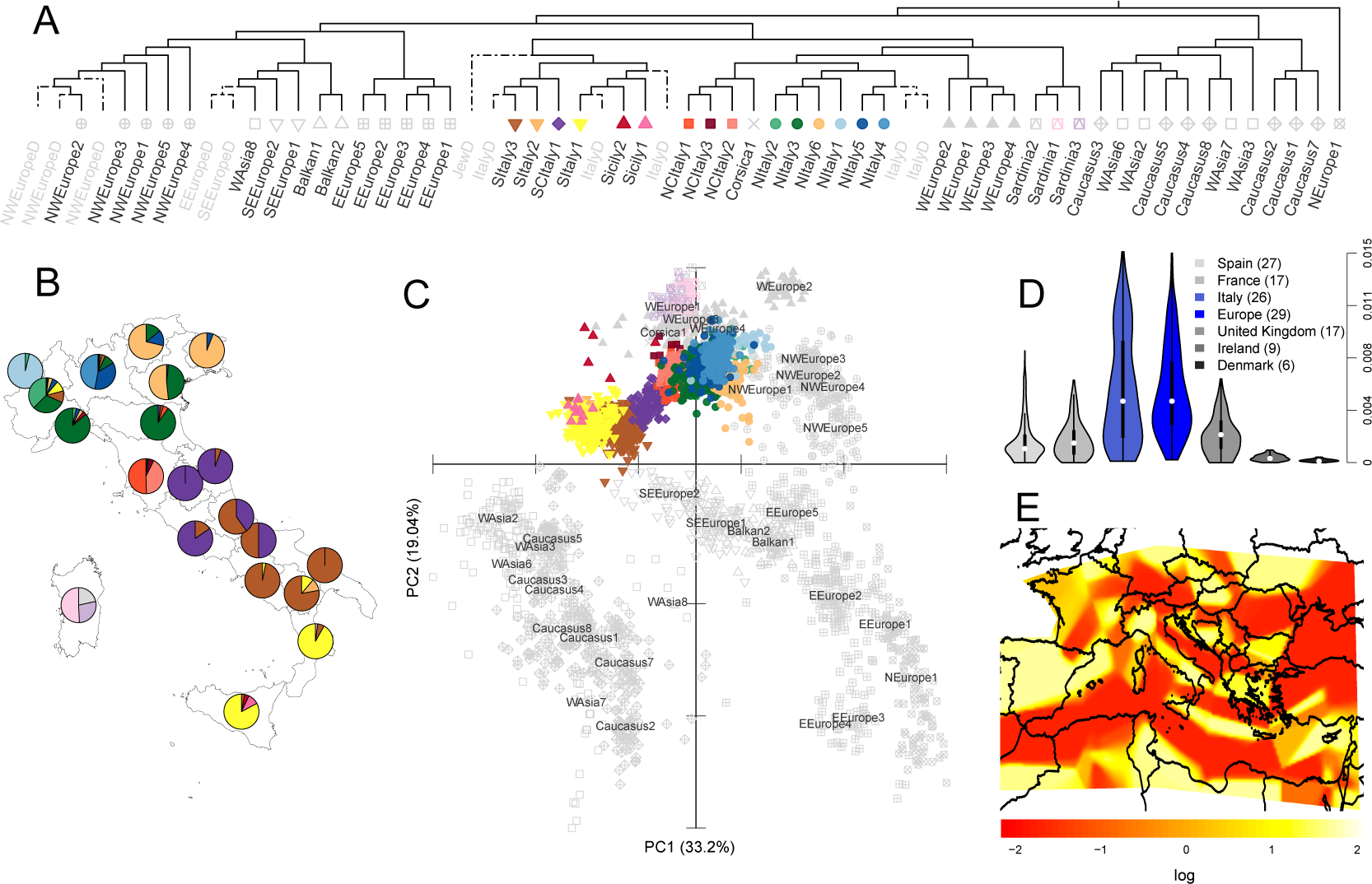
Genetic structure of the Italian populations. A) Simplified dendrogram of 3,057 Eurasian samples clustered by the fS algorithm using the CP output (complete dendrogram in fig. S2A); each leaf represents a cluster of individuals with similar copying vectors; clusters with more than five individuals are labelled in black; Italian clusters are colour coded; grey labels ending with the D letter refer to clusters containing less than five individuals or individuals of uncertain origin that have been removed in the following analyses. B) Pie charts summarizing the relative proportions of inferred fS genetic clusters for all the 20 Italian administrative regions (colours as in A). C) PCA based on CP chunkcount matrix (colours as in A); the centroid of the individuals belonging to non-Italian clusters is identified by the label for each cluster. D) Between-clusters F_st_ estimates within European groups; clusters were generated using only individuals belonging to the population analysed (Materials and Methods, Supplementary materials); the number of genetic clusters analysed for each population is reported within brackets; for the comparisons across Europe, the cluster NEurope1 containing almost exclusively Finnish individuals was excluded (F_st_ estimates for Italian and European clusters are in data file S3); F_st_ distributions statistically different from the Italian set are in grey. E) Estimated Effective Migration Surfaces (EEMS) analysis in Southern Europe; colours represent the log10 scale of the effective migration rate, from low (red) to high (yellow).

Italian clusters separated into three main groups: Sardinia, Northern (North/Central-North Italy) and Southern Italy (South/Central-South Italy and Sicily); the former two were close to populations originally from Western Europe, while the latter was in proximity of Middle East groups (Fig. 1A, fig. S2, data file S2). The cluster-composition of the administrative regions of Italy provided further evidence for geographic structuring (Fig. 1B) with the separation between Northern and Southern areas being shifted North along the peninsula; the affinity to Western and Middle Eastern populations was also evident in the haplotype-based PCA (Fig. 1C), allele frequency PCA (fig. S3) and the ADMIXTURE analysis (fig. S4).

These observations were replicated using a subset of the dataset genotyped for a larger number of SNPs (High Density Dataset, HDD, including 591,217 SNPs; see Materials and Methods, Supplementary materials, Fig. 1B, table S1). Recent migrants and admixed individuals, as identified on the basis of their copying vectors (fig. S5, fig. S6, table S2), were removed in subsequent CP/fS analyses (see Supplementary materials).

We explored the degree of within-country differentiation by comparing the distribution of FST values among fS genetic clusters in Italy with the ones in several European countries (*13*–*16*) and across the whole of Europe. Clusters within Italy were significantly more different from each other than within any other country here included (median Italy: 0.004, data file S3; range medians for listed countries 0.0001-0.002) and showed differences comparable with estimates across European clusters (median European clusters: 0.004, Fig. 1D, see Materials and Methods, Supplementary materials). The analysis of the migration surfaces (EEMS) (*17*) highlighted several barriers to gene flow within and around Italy but also suggested the existence of migration corridors in the southern part of the Adriatic and Ionian Sea, and between Sardinia, Corsica and continental Italy (Fig. 1E; fig. S7) (*11*).

### Multiple ancient ancestries in Italian clusters

We investigated the ancestry composition of modern clusters by testing different combination of ancient samples using the CP/NNLS pipeline, a previously implemented analysis that reconstructs the profiles of modern populations as the combination of the “painted” profiles of different ancient samples by using a “mixture fit” approach based on a non-negative least square algorithm (NNLS) (*13*, *18*, *19*). We applied this approach to ancient samples using the unlinked mode implemented in CP, similarly to other routinely performed analyses based on unlinked markers or allele frequency, such as qpAdm and ADMIXTURE. In addition, data from modern individuals (FMD) were harnessed as donor populations (see Materials and Methods, Supplementary materials). Following Lazaridis et. al 2017 (*10*), we performed two separate CP/NNLS analyses, *“Ultimate”* and *“Proximate”*, referring to the least and the most recent putative sources, respectively (Fig. 2, fig. S8, fig. S9). In the *Ultimate* analysis, all the Italian clusters were characterised by relatively high amounts of Anatolian Neolithic (AN), ranging between 56% (SItaly1) and 72% (NItaly4), distributed along a North-South cline (Spearman ρ = 0.52, p-value < 0.05; Fig. 2A-C, fig. S8A), with Sardinians showing values above 80%. A closer affinity of Northern Italian than Southern Italian clusters to AN was also supported by D-statistics (fig. S10). The remaining ancestry was mainly assigned to WHG (Western Hunter-Gatherer), CHG and EHG. In particular, the first two components were more present in populations from the South (higher estimates in SItaly1 ∼13% and SItaly3 ∼ 24% for WHG and CHG respectively), while the latter was more common in Northern clusters (NItaly6 = 15%). These observations suggest the existence of different secondary sources contributions to the two edges of the peninsulas, with the North affected more by EHG-related populations and the South affected more by CHG-related groups. Iran Neolithic (IN) ancestry was detected in Europe only in Southern Italy.

**Fig. 2.**
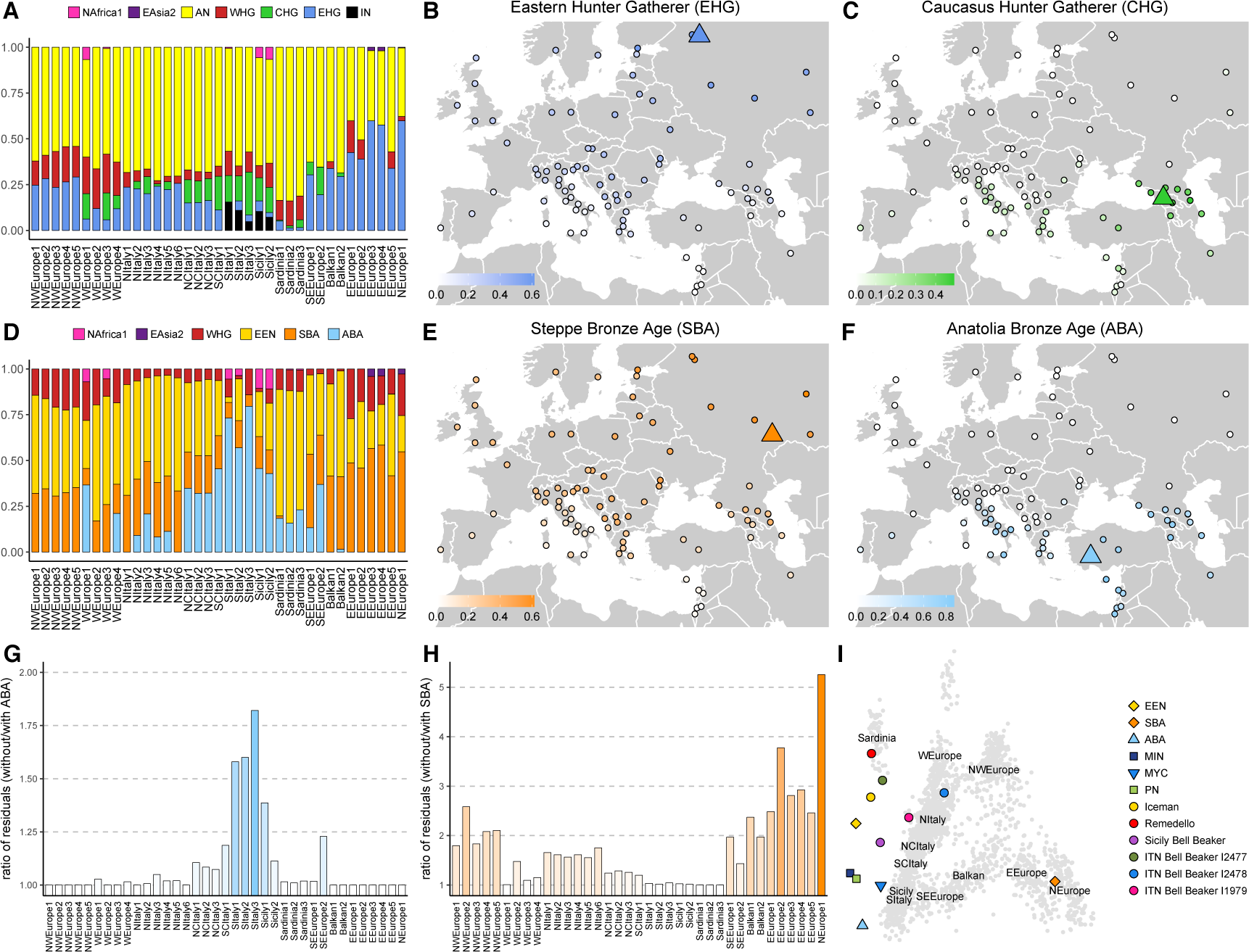
Ancient ancestries in Western Eurasian modern-day clusters and Italian ancient samples. A, D) CP/NNLS analysis on all Italian and European clusters using as donors different sets of ancient samples and two modern clusters (NAfrica1: North Africa, EAsia2: East Asia) (full results in fig. S8). A) *Ultimate sources:* AN, Anatolian Neolithic (Bar8); WHG, Western Hunter Gatherer (Bichon); CHG, Caucasus Hunter Gatherer (KK1); EHG, Eastern Hunter Gatherer (I0061); IN, Iranian Neolithic (WC1). B) EHG and C) CHG ancestry contributions in Western Eurasia, as inferred in A and fig. S8A (Supplementary materials). D) Same as in A, using *Proximate* sources: WHG, Western Hunter Gatherer (Bichon); EEN, European Early Neolithic (Stuttgart); SBA, Bronze Age from Steppe (I0231); ABA, Bronze Age from Anatolia (I2683). E) SBA and F) ABA ancestry contributions, as inferred in D and fig. S8B. Triangles refer to the location of ancient samples used as sources (see data file S1). G): ratio of the residuals in the NNLS analysis (Materials and Methods, Supplementary materials) for all the Italian and European clusters when ABA was excluded and included in the set of *Proximate* sources; H) as in G), but excluding/including SBA instead of ABA; J) Ancient Italian and other selected ancient samples projected on the components inferred from modern European individuals. Labels are placed at the centroid of the individuals belonging to the indicated clusters.

North-South differences across Italy were also detected in the *Proximate* analysis. When *Proximate* sources were evaluated, SBA contribution ranged between 33% in the North and 6% in the South of Italy, while ABA (Anatolia Bronze Age) showed an opposite distribution (Fig. 2D-F, fig. S9), in line with the results based on the D statistics (fig. S10, fig. S11), and mirroring the EHG and CHG patterns, respectively. Contrary to previous reports, the occurrence of CHG as detected by the CP/NNLS analysis did not mirror the presence of Steppe Bronze Age (SBA), with several populations testing positive for the latter but not for the former ((*6*), Fig. 2, fig. S8). We therefore speculate that our approach might in general underestimate the presence of CHG across the continent; however, we note that even considering this scenario, the excess of Caucasus related ancestry detected in the South of the European continent, and in Southern Italy in particular, is striking and unexplained by currently proposed models for the peopling of the continent.

Interestingly, clusters belonging to the North had more EEN (European Early Neolithic) than Southern ones, which in turn were composed by an higher fraction of ABA, although the high AN-related component in both these ancient groups might have affected the exact source identification. The relevance of ABA in Italy was additionally supported by the reduced fit of the NNLS (sum of the squared residuals; Materials and Methods, Supplementary materials) when the *Proximate* analysis was run excluding ABA. Results were similar to the full *Proximate* analysis for most of the European clusters, but not for Southern European groups, where the residuals were almost up to twice as much when ABA was not included as a source (Fig. 2G). A similar behaviour, but for Northern Italian and most of the European clusters, was observed when SBA was removed from the panel of *Proximate* sources (Fig. 2H). The closer affinity of the Southern Italian clusters to ABA was also highlighted by the PCA and ADMIXTURE analysis on ancient and modern samples (Fig. 2I, fig. S12, fig. S13, fig. S14) and significantly higher ABA ancestry in Southern than Northern Italy, as estimated by NNLS analysis (Fig. 2D, Student’s t-Test p-value < 0.05, Supplementary materials). We also noted that in the Balkan peninsula signatures related to ABA were present but less evident than in Southern Italy across modern-day populations, possibly masked by historical contributions from Central Europe (*20*, *21*) (Fig. 2, Fig. 3, fig. S8B). Overall, SBA and ABA appear to have very different distribution patterns in Europe: continent-wide the former, more localised (in the South) the latter. Similar results were obtained when other Southern European ancient sources replaced ABA in the Proximate analysis (fig. S9, Materials and Methods, Supplementary materials). These results were confirmed by qpAdm analysis. When two sources were evaluated, a large AN contribution was supported only in one cluster (SItaly2), while the vast majority of supported models included ABA, Minoan or Mycenaean and one of the hunter-gatherer groups or SBA (table S3, table S4). When three possible sources were allowed, AN was supported for all the Southern Italian clusters, mostly in association with EHG/WHG/SBA and CHG/IN. Nevertheless, all the analysed clusters, could be modelled as a combination of ABA, SBA and European Middle-Neolithic/Chalcolithic, their contributions mirroring the pattern observed in the CP/NNLS analysis (fig. S15, table S3, table S4). North African contributions, ranging between 3.8% (SCItaly1) to 14.5% (SItaly1) became evident when combinations of five sources were tested. Sardinian clusters were consistently modeled as AN+WHG+CHG/IN across runs, with the inclusion of North Africa and SBA when different number of sources were considered. The qpAdm analyses of Italian HDD clusters generated similar results (Materials and Methods, Supplementary materials, table S4). In order to obtain insights about the relationship between ancient and modern groups, we performed the same qpAdm analysis on post-Neolithic/Bronze Age Italian individuals (fig. S15, table S5). Iceman and Remedello, the oldest Italian samples here included (3,400-2,800 BCE, Before Current Era), were composed by high proportions of AN (74 and 85%, respectively). The Bell Beaker samples of Northern Italy (2,200-1,930 BCE) were modelled as ABA and AN + SBA and WHG, although ABA was characterised by large standard errors but the detection of Steppe ancestry, at 14%, was more robust. On the other hand Bell Beaker samples from Sicily (2,500-1,900 BCE) were modelled almost exclusively as ABA, with less than 5% SBA. Despite the fact that the small number of SNPs and prehistoric individuals tested prevents the formulation of conclusive results, differences in the occurrence of AN ancestry, and possibly also Bronze Age related contributions, are suggested to be present between ancient samples from North and South Italy. Differences across ancient Italian samples were also supported by their projections on the PCA of modern-day data (Fig. 2I). Remedello and Iceman clustered with European Early Neolithic samples, together with one of the three Bell Beaker individuals from North Italy, as previously reported (*22*), and modern-day Sardinians. The other two Bronze Age North Italian samples clustered with modern North Italians, while the Bell Beaker sample from Sicily was projected in between European Early Neolithic, Bronze Age Southern European and modern-day Italian samples (Fig. 2I).

**Fig. 3.**
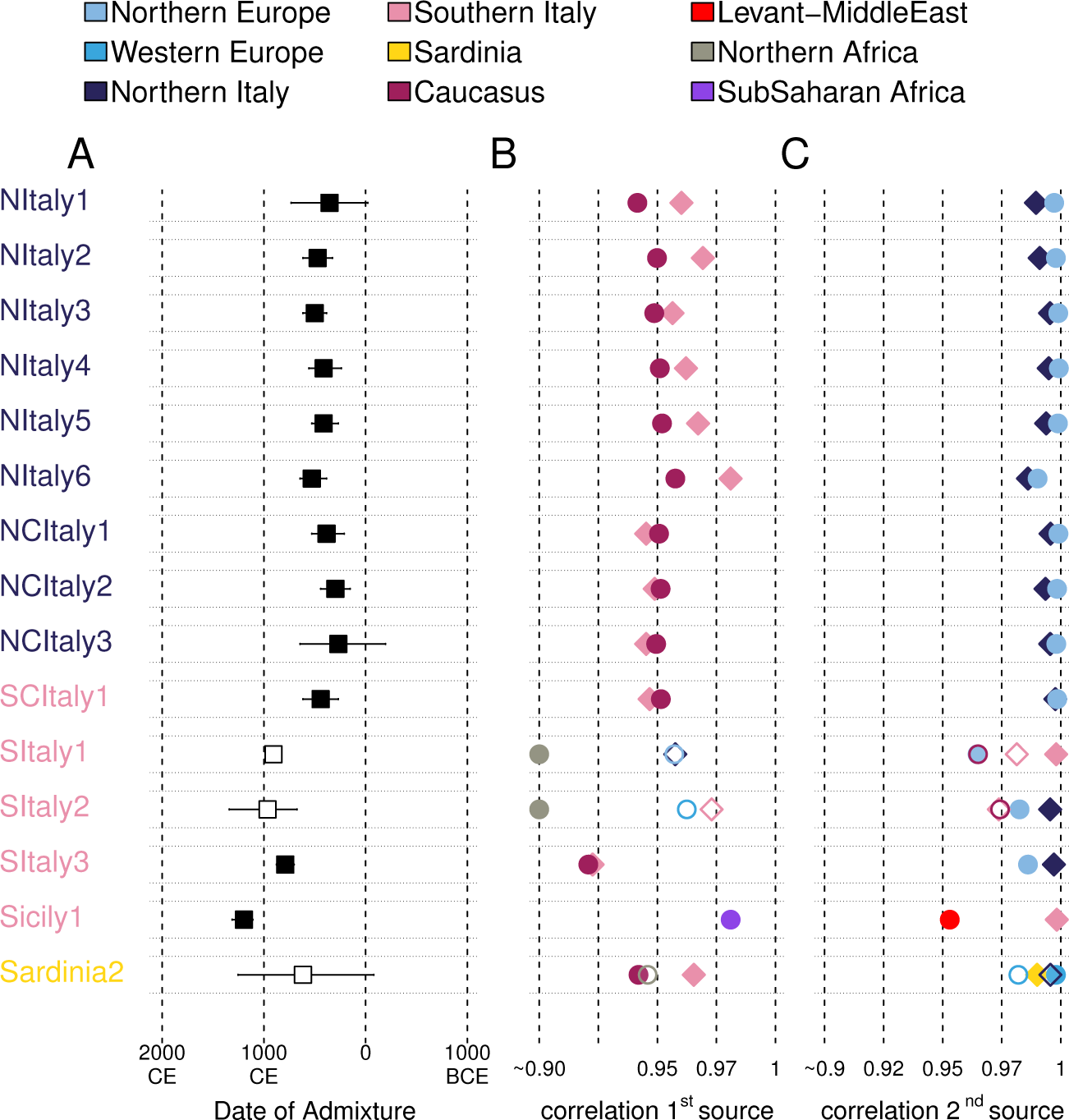
Admixture events inferred by GLOBETROTTER (GT). A) Dates of the events inferred in the GT “noItaly” analysis on all the Italian clusters (els as in Fig. 1A and data file S2; full results in fig. S16 and table S7; see Materials and Methods, Supplementary materials); lines encompassed the 95% CI. GT events were distinguished in “one date” (black squares; 1D in table S7) and “one date multiway” (white squares; 1MW). B) Correlation values between copying vectors of 1 ^st^ source(s) identified by GT and the best proxy in the noItaly analysis (circles) or the best proxy among Italian clusters (diamonds). C) Same as in B, referring to 2 ^nd^ source(s) copying vectors. Empty symbols refer to additional 1 ^st^ (B) and 2 ^nd^ (C) sources detected in multiway events. African best proxies in (B) for clusters SItaly1 and SItaly2 were plotted on the 0.90 boundary for visualisation only, the correlation values being 0.78 and 0.87 respectively. Colours of symbols refer to the ancestry to which proxies were assigned (see Materials and Methods, Supplementary materials).

### Historical admixture

In order to investigate the role of historical admixture events in shaping the modern distribution of ancient ancestries, we generated the admixture profiles of Italian and European populations using GLOBETROTTER (GT, (*21*)) (Fig. 3, fig. S16, table S6, table S7).

We discussed here the results based on the full modern dataset (FMD) as it provided a wider coverage at population level.

We run the analysis excluding the Italians as donors in order to reduce copying between highly similar groups (GT “noItaly” analysis; Fig. 3). The events detected in Italy occurred mostly between 1,000 and 2,000 years ago (ya), and extended to 2,500ya in the rest of Europe (Fig. 3A and fig. S16). Clusters from Caucasus and North-West Europe were identified all across Italy as best-proxies for the admixing sources, while Middle Eastern and African clusters were identified as best proxies only in Southern Italian clusters and Sardinia (Fig. 3B, C). We noted that when we extended the search for the best-proxies to include also Italian clusters, these were as good as or better proxies than clusters from the Caucasus and the Middle East. On the other hand, North-West European and African clusters were usually still better proxies than groups from any other area (Fig. 3B, C). Notably, Eastern and Middle Eastern clusters were not detected as best proxies when we run the GT analysis including all clusters as donors, contrary to African, European and Italian groups (“GTall” analysis; table S6). Overall these results supported a scenario in which gene flow mostly occurred between resident Italian sources and non-Italian sources. SBA and ABA ancestries were detected in Italian and non-Italian best-proxies (Fig. 2D, Fig. 3, table S6, table S7), which suggests that part of these ancestries arrived from outside Italy in historical times, but also that these components were already present in Italian groups at the time of these admixture events. Episodes of gene flow were also detected in Sardinia, combining signals from both the African continent and North West Europe. MALDER results for the more recent episodes replicated the admixture pattern identified by GT (fig. S16, table S8).

### The Neanderthal legacy across Italy and Europe

The variation in ancestry composition reported across Italy and Europe is expected to influence other aspects of the genetic profiles of European populations, including the presence of archaic genetic material (*6*). We investigated the degree of Neanderthal ancestry in Italian and other Eurasian populations by focusing on SNPs tagging Neanderthal introgressed regions (*23*, *24*). SNPs were pruned for LD and a final set of 3,969 SNPs was used to estimate the number of Neanderthal alleles in samples genotyped for the Infinium Omni2.5-8 Illumina beadchip. Asian and Northern European populations had significantly more Neanderthal alleles than European and Southern European groups respectively, as previously reported (*25*–*28*), with significant differences also highlighted within Italy (Fig. 4A, B). Contributions from African groups possibly influenced these patterns, particularly in Southern European populations (*20*) (Fig. 2, Fig. 3). However differences within Europe and Italy were still present once individuals belonging to clusters with African contributions were removed (fig. S17, see Materials and Methods, Supplementary methods). Ancient samples have been reported to differ in the amount of Neanderthal DNA due to variation in the presence of a so-called “Basal Eurasian” lineage, stemming from non-Africans before the separation of Eurasian groups and harbouring only a negligible fraction of Neanderthal ancestry (*6*). Consistent with this (*6*), we found the estimated amounts of Basal Eurasian and Neanderthal to be negatively correlated across modern day European clusters (Fig. 4C, fig. S18, fig. S19), irrespective of the removal of all the clusters admixed with African sources (see Materials and Methods, Supplementary materials; fig. S17).

**Fig. 4.**
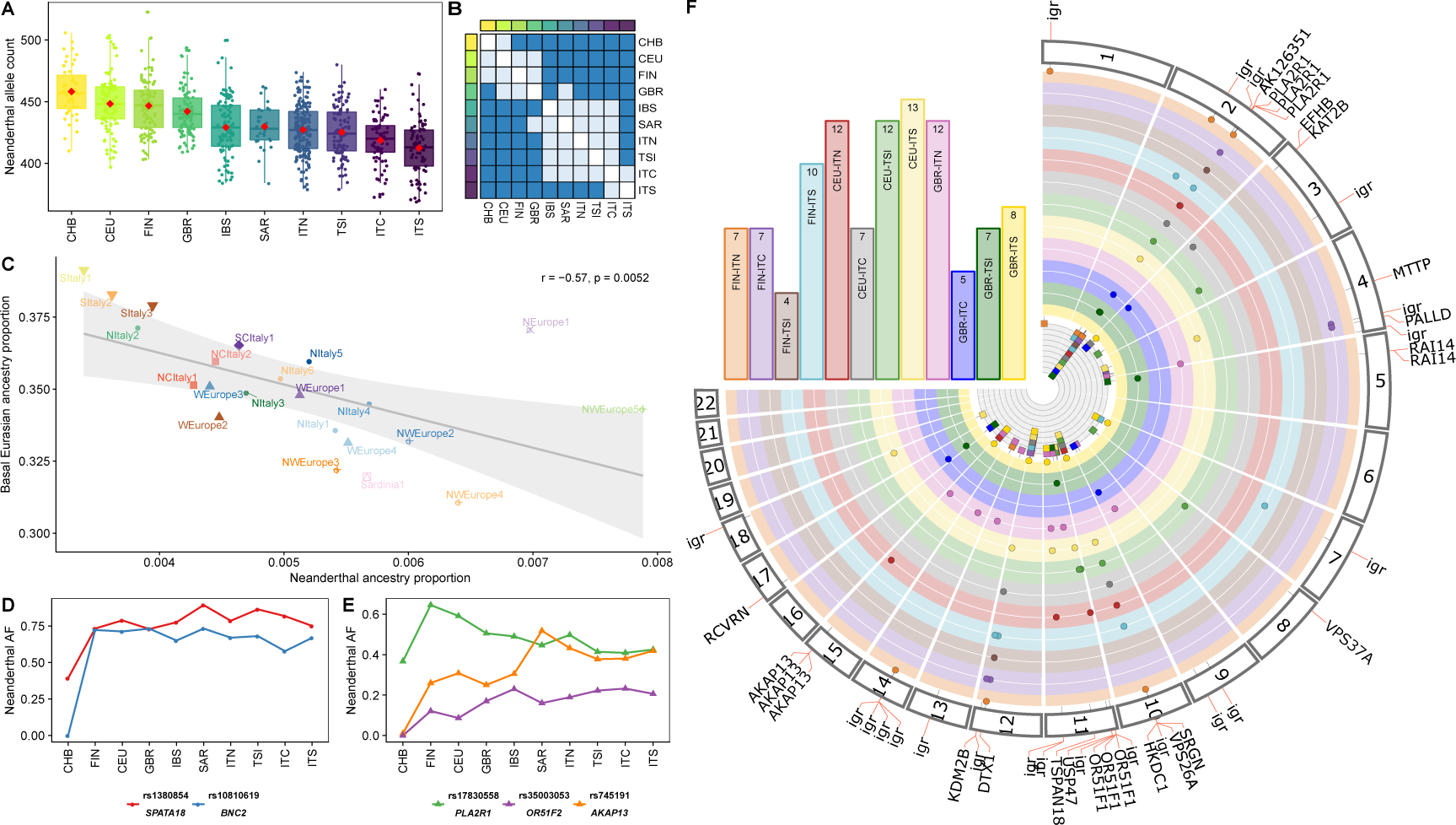
Neanderthal ancestry distribution in Eurasian populations. A) Neanderthal allele counts in individuals from Eurasian populations, sorted by median values on 3,969 LD-pruned Neanderthal tag-SNPs. CEU, Utah Residents with Northern and Western European ancestry; GBR, British in England and Scotland; FIN, Finnish in Finland; IBS, Iberian Population in Spain; TSI, Tuscans from Italy; ITN, Italians from North Italy; ITC, Italians from Central Italy; ITS, Italians from South Italy; SAR, Italians from Sardinia; CHB, Han Chinese. B) Matrix of significances based on Wilcoxon rank sum test between pairs of populations including (lower triangular matrix) and removing (upper) outliers (Materials and Methods, Supplementary materials; dark blue: adj p-value < 0:05; light blue: adj p-value > 0:05). C) Correlation between Neanderthal ancestry proportions and the amount of Basal Eurasian ancestry in European clusters (Materials and Methods, Supplementary materials). D, E) Neanderthal allele frequency (AF) for selected SNPs within the indicated genes: D) high frequency alleles in Europe; E) North-South Europe divergent alleles. F) Comparisons between Northern European and Italian populations (excluding Sardinia). Bars refer to comparison for reported pairs of populations; the number of NTT SNPs is reported within bars. Each section of the circos represents a tested chromosome; points refer to NTT SNPs. Colours, same as for bars; igr: intergenic region variant.

The variation in Neanderthal ancestry was also reflected at specific loci. A total of 144 SNPs were identified among the Neanderthal-tag SNPs showing the largest differences in allelic frequency in genome-wide comparisons across Eurasian and African populations (see Materials and Methods, Supplementary materials - Neanderthal-Tag SNPs within the Top 1% of the genome-wide distributions of each of the 55 pairwise population comparisons - NTT SNPs; fig. S20). The top 1% of each distribution was significantly depleted in Neanderthal SNPs (see Materials and Methods, Supplementary materials, table S9), in agreement with a scenario of Neanderthal mildly deleterious variants being removed more efficiently in human populations (*29*–*31*).

The 50 genes containing NTT SNPs were enriched for phenotypes related to facial morphology, body size, metabolism and muscular diseases (see Materials and Methods, Supplementary materials, data file S4). A total of 34 NTT SNPs were found to have at least one known phenotypic association (*32, 33*) (data file S4). Among these, we found Neanderthal alleles associated with increased gene expression in testis and in skin after sun exposure (SNPs within the *IP6K3* and *ITPR3* genes), susceptibility to cardiovascular and renal conditions (*AGTR1*), and Brittle cornea syndrome (*PRDM5*) (*24*). NTT SNPs between European and Asian/African populations included previously reported variants in *BNC2* and *SPATA18* genes (*23*, *34*, *35*) (see Materials and Methods, Supplementary materials, Fig. 4D), while 80 NTT SNPs were involved in at least one comparison between Northern (CEU, GBR and FIN) and Southern European populations (IBS and Italian groups). Among these SNPs, three mapped to the Neanderthal introgressed haplotype hosting the *PLA2R1* gene, the archaic allele at these positions reaching frequencies of at least 43% in Northern European and at most of 35% in Southern European populations (Fig. 4E, F). Ten SNPs showed an opposite frequency gradient: seven mapped to one Neanderthal introgressed region spanning the *OR51F1, OR51F2* and *OR52R1* genes (Fig. 4E, F), and the other three identified regions hosting the *AKAP13* gene, within one of the high frequency European Neanderthal introgressed haplotypes recently reported (*36*) (Fig. 4E, F).

## Discussion

The pattern of variation reported across Italian groups appears geographically structured in three main regions: Southern and Northern Italy and Sardinia. The North-South division in particular appeared as shaped by the distribution of Bronze Age ancestries with signatures of different continental hunter-gatherer groups. The results of the analyses of both modern and ancient data suggest that ancestries related to Caucasus and Eastern hunter-gatherers were possibly initially brought in Italy by at least two different contributions from the East. Of these, one is the well-characterised SBA signature ultimately associated with the nomadic groups from the Pontic-Caspian steppes. This component entered Italy from mainland Europe and was present in the peninsula in the Bronze Age, as suggested by its presence in Bell Beaker samples from North Italy (table S5). SBA ancestry continued to arrive from the continent up until historical times (Fig. 3). The other contribution is ultimately associated with CHG ancestry and affected predominantly the South of Italy, where it now represents a substantial component of the ancestry profile of local populations. This signature is still uncharacterised in terms of precise dates and origin; however such ancestry was possibly already present during the Bronze Age in Southern Italy (table S5) and was further supplemented by historical events (Fig. 3).

The very low presence of CHG signatures in Sardinia and in older Italian samples (Remedello and Iceman) but the occurrence in modern-day Southern Italians might be explained by different scenarios, not mutually exclusive: 1) population structure among early foraging groups across Italy, reflecting different affinities to CHG; 2) the presence in Italy of different Neolithic contributions, characterised by different proportion of CHG-related ancestry; 3) the combination of a post-Neolithic, prehistoric CHG-enriched contribution with a previous AN-related Neolithic layer; 4) A substantial historical contribution from Southern East Europe across the whole of Southern Italy. No substantial structure has been highlighted so far in pre-Neolithic Italian samples (*8*). An arrival of the CHG-related component in Southern Italy from the Southern part of the Balkan Peninsula is compatible with the identification of genetic corridors linking the two regions (Figure 1E, (*11*)) and the presence of Southern European ancient signatures in Italy (Figure 2). The temporal appearance of CHG signatures in Anatolia and Southern East Europe in the Late Neolithic/Bronze Age suggests its relevance for post-Neolithic contributions (*37*). Additional analyses of aDNA samples from around this time in Italy are expected to clarify what scenario might be best supported.

Historical events possibly involving continental groups at the end of Roman Empire and African contributions following the establishment of Arab kingdoms in Europe around 1,000 ya (*20*, *21, 38*–*40*) played a role in further shaping the ancestry profiles of the Italian populations.

Despite Sardinia was confirmed as being the most closely related population to Early European Neolithic farmers (Figure 2D, I), there is no evidence for a simple genetic continuity between the two groups. Sardinia, and the rest of Italy, experienced in fact historical episodes of gene-flow (*4*) (Fig. 2, Fig. 3, table S3, table S4) that contributed to the further dispersal of ancient ancestries and the introduction of other components, including African ones.

It has been previously reported that variation in the effective population size might explain differences in the amount of Neanderthal DNA detected in European and Asian populations (*24, 27*, *41*). Additional Neanderthal introgression events in Asia and gene-flow from populations with lower Neanderthal ancestry in Europe possibly provide further explanations for differences in Neanderthal occurrence across populations (*42*). The spatial heterogeneity of Neanderthal legacy within Europe here reported appears as the result of ancient and historical events which brought together in different combinations groups harbouring different amounts of Neanderthal genetic material. While these events have shaped the overall continental distribution of Neanderthal DNA, locus-specific differences in the occurrence of Neanderthal alleles are also expected to reflect selective pressures acting on these variants since their introgression in the populations (*30, 31*).

The variation in ancestry composition detected across Italy extends to neighbour regions and appears to combine historical contributions and ancient stratification. The differences between Northern and Southern Italian populations are possibly reflecting long-term differential links with Central and Southern Europe respectively, with additional contributions from the African continent for the Southern part of Italy and Sardinia.

The multifaceted admixture profile here sketched provides an interpretative framework for the processes that have shaped Southern European genetic variation. The inclusion of ancient samples spanning diachronic and geographic transects from the Italian peninsula and nearby regions will help in clearing up further questions about the temporal and spatial dynamics of these processes.

## Materials and Methods

### Analysis of modern samples

#### Dataset

Two hundred and twenty-four samples are here present for the first time. Of these, 167 Italians and 6 Albanians were specifically selected and sequenced for this project with two versions (1.2 and 1.3) of the Infinium Omni2.5-8 Illumina beadchip, while 57 additional Italians and Europeans were previously sequenced with Illumina 660W and are presented here for the first time (Supplementary materials, table S1). Two separate world-wide datasets were prepared. The Full Modern Dataset (FMD) included 4,852 samples (1,589 Italians) and 218,725 SNPs genotyped with Illumina arrays; the High Density Dataset (HDD) contained 1,651 samples (524 Italians) and 591,217 SNPs genotyped with the Illumina Omni array (Supplementary materials).

The merging, the removal of ambiguous C/G and A/T and triallelic markers, the exclusion of related individuals and the discarding of SNPs in linkage disequilibrium (LD) were performed using PLINK1.9 (*43, 44*). Only autosomal markers were considered.

#### Haplotype analysis (CHROMOPAINTER, CP, and fineSTRUCTURE, fs)

Phased haplotypes were generated using SHAPEIT(*45*) and applying the HapMap b37 genetic map.

CP was employed to generate a matrix of recipient individuals “painted” as a combination of donor samples (copying vector). Three runs of CP were done for each dataset generating three different outputs: (i) a matrix of all the individuals “painted” as a combination of all the individuals, for cluster identification and GT analysis; (ii) a matrix of all Italians as a combination of all Italians, for FST analysis; (iii) a matrix of all the samples as a combination of all the other samples but excluding Italians, for “local” GT analysis.

Clusters were inferred using fineSTRUCTURE (fS). After an initial search based on the “greedy” mode, the dendrogram was processed by visual inspection (*18*, *20*) according to the geographical origin of the samples. The robustness of the cluster was obtained by processing the MCMC pairwise coincidence matrix (Supplementary materials).

#### Cluster Self-Copy Analysis

Recently admixed individuals were identified as those copying from members of the cluster they belong less than the amount of cluster self-copying for samples with all the four grandparents from the same geographic region (Supplementary materials).

#### Principal Component Analysis (PCA)

PCA was performed on CP chunkcount matrix (Supplementary materials) and was generated using the prcomp() function on R software (*46*). Allele frequencies PCA was performed using smartpca implemented in the EIGENSOFT(*47*) after pruning the datasets for LD.

#### Characterization of the migration landscape (EEMS analysis)

Estimated Effective Migration Surfaces analysis (EEMS) (*17*) was performed estimating the average pairwise distances between population using bed2diffs tool and the resulting output was visualised by using the Reems package (*17*).

#### ADMIXTURE analysis

ADMIXTURE1.3.0 software (*48*) was used performing 10 different runs using a random seed. The results were combined with CLUMPP (*49*) using the largeKGreedy algorithm and random input orders with 10,000 repeats. *Distruct* implemented in CLUMPAK (*50, 51*) was then used to identify the best alignment of CLUMPP results. Results were processed using R statistical software (*46*).

#### FST estimates among clusters

Pairwise FST estimates among newly generated Italian clusters and among originally generated European clusters (Supplementary materials) were inferred using smartpca software implemented in the EINGESOFT package (*47*). Comparisons between the FST distributions were performed using a Wilcoxon rank sum test in R programming language environment.

#### The time and the sources of admixture events (GT analysis and MALDER analysis)

Times of haplotype-dense data admixture events were investigated using GLOBETROTTERv2 software. GT was employed using two approaches: complete and non-local (referred as “noItalian”, Supplementary materials), in default modality (*13, 20, 52*). The difference between the two approaches was the inclusion or the exclusion respectively of all the Italian clusters as donors in the CP matrix used as input file. To improve the precision of the admixture signals, “null.ind 1” parameter was set (*52*). Unclear signals were corrected using the default parameters and a total of 100 bootstraps were performed. MALDER uses allele frequencies to dissect the time of admixture signals. The best amplitude was identified and used to calculate a Z-score (Supplementary materials). A Z-score equal or lower than 2 identifies not significantly different amplitude curves (*53, 54*) (Supplementary materials).

Sources for both GT and MALDER were grouped in different ancestries as indicated in the legend of Fig. 3, fig. S16.

The expression (1950 – (g +1)* 29), where g is the number of generation, was used to convert into years the GT and MALDER results, negative numbers were preceded by BCE (Before Current Era) letters.

### Analyses including ancient samples

#### Dataset

In order to explore the extent to which the European and Italian genetic variation has been shaped by ancient demographic events, we merged modern samples from FMD with 63 ancient samples selected from recent studies (*6, 7, 10, 22*, *37, 55*–*57*) (data file S1).

#### Principal Component Analysis (PCA)

We performed two principal components analyses with the EIGENSOFT (*47*) smartpca software and the *“lsqproject”* and *“shrinkmode”* option, projecting the ancient samples on the components inferred from modern European, West Asian and Caucasian individuals and, then, only on modern European clusters. In order to evaluate the potential impact of DNA damage in calling variants from aDNA samples, we repeated the PCA with the 63 ancient samples and modern European, Caucasian and West Asian samples by removing transition polymorphisms and recorded significant correlations for the localisation of ancient samples along PC1 and PC2 (r > 0.99, p-value < 0.05).

#### ADMIXTURE analysis

We projected the ancient samples on the previously inferred ancestral allele frequencies from 10 ADMIXTURE (*48*) runs on modern samples (see “Analysis of modern samples” section and Supplementary materials). We used CLUMPP(*49*) for merging the resulting matrices and *distruct (51*) for the visualization.

#### D-STATISTICS

We tested for admixture using the D-statistics as implemented in the qpDstat tool in the software ADMIXTOOLS v4.2 (*58*). We performed the D-statistic analyses evaluating the relationship of Italian cluster with AN, ABA and SBA. In details, we performed the the D-statistics D(Ita1,Ita2,AN/ABA/SBA,Mbuti) where Ita1 and Ita2 are the different clusters composed mainly by italian individuals as inferred by fineStructure.

#### CHROMOPAINTER (CP)/Non-Negative Least Squares (NNLS) analysis

We used an approach based on the software CP (*12*, *59*) and a slight adaptation of the non-negative least square (NNLS) function (*13*, *18*, *19*) to estimate the proportions of the genetic contributions from ancient population to our modern clusters. We run CP using the “unlinked” mode (*55*) and the same Ne and θ parameters of the modern dataset and we painted both modern and ancient individuals, using only modern samples as donors (*55, 56*). Then we “inverted” the output of CP by solving an appropriately formulated NNLS problem, producing a painting of the modern clusters in terms of the ancients. We applied this combined approach on different sets of ancient samples (*Ultimate* and various combinations of *Proximate* sources).

The goodness of fit of the NNLS was measured evaluating the residuals of the NNLS analysis. In details, we focused on the Proximate sources, and compared the sum of squared residuals when ABA or SBA were included/excluded as putative sources.

#### qpAdm analysis

We used the ancestral reconstruction method qpAdm, which harnesses different relationships of populations related to a set of outgroups (**eg.** f4[Target, O1, O2, O3]).

In details, for each tested cluster of the FMD and HDD, we have evaluated all the possible combinations of N “left” sources with N={2‥5}, and one set of right/left Outgroups (Supplementary materials).

For each of the tested combinations we used qpWave to evaluate if the set of chosen outgroups is able to I) discriminate the combinations of sources and II) if the target may be explained by the sources. We used a p-value threshold of 0.01. Finally, we used qpAdm to infer the admixture proportions and reported it and the associated standard errors in Supplementary table S3 and table S4. In addition, we performed the same analysis for Iceman, Remedello and Bell Beaker individuals from Sicily and North Italy (table S5).

### Archaic contribution

#### Dataset

We assembled an additional high density dataset by retaining only samples genotyped on the Illumina Infinium Omni2.5-8 BeadChip from our larger modern dataset. In particular, we included seven populations from the 1000 Genomes Project: the five European populations (Northern European from Utah - CEU, England - GBR, Finland - FIN, Spain - IBS, Italy from Tuscany - TSI), one from Asia (Han Chinese - CHB) and one from Africa (Yoruba from Nigeria - YRI). We also retained 466 Italian samples, whose four grandparents were born in the same Italian region. The Italian samples were broadly clustered according to their geographical origin into Northern (ITN), Central (ITC), Southern (ITS) Italians and Sardinians (SAR), while TSI samples from 1000 Genome Project formed a separate cluster (table S10).

From this dataset, we extracted 7,164 Neanderthal SNPs tagging Neanderthal introgressed regions (*24*). In order to select which allele was inherited from Neanderthals, we chose the one from the Altai Neanderthal (*41*) genome when it was homozygous and the minor allele in YRI when it was heterozygous.

#### Number of Neanderthal alleles in present-day human populations

After pruning variants in linkage disequilibrium, we counted the number of Neanderthal alleles considering all the tag-SNP across all samples. Then, we compared the distribution of Neanderthal allele counts across populations with the two-sample Wilcoxon rank sum test. We repeated the same analyses after removing outlier individuals.

#### Basal Eurasian ancestry and Neanderthal contribution

In order to infer the proportion of Basal Eurasian present in European populations (*6, 7*), we used the f4 ratio implemented in the ADMIXTOOLS package (*58*) in the form f4(Target, Loschbour, Ust_Ishim, Kostenki14)/ f4(Mbuti, Loschbour, Ust_Ishim, Kostenki14). We repeated this approach to infer the Neanderthal ancestry, in the form f4 (Mbuti, Chimp Target, Altai)/ f4(Mbuti, Chimp, Dinka, Altai) (fig. S18, fig. S19). We then performed the same analyses by grouping the modern individuals according to the CP/fS inferred clusters (“Analysis of modern samples” section) and retained only clusters with at least 10 samples (Fig. 4)

#### African ancestry and Neanderthal legacy

The impact of African contributions in shaping the amount of Neanderthal occurrence was evaluated by exploring how the removal of the clusters showing African gene-flow as detected by GT analysis (Fig. 3) and how individuals belonging to these clusters affected the correlation between Basal Eurasian/Neanderthal estimates and the degree of population differentiation in the amount of Neanderthal alleles, respectively (Supplementary materials; fig. S17).

#### Comparison of Neanderthal allele frequencies across modern populations

We computed the allele frequency differences for every SNPs for each of the possible pairs of the eleven populations in our dataset, thus obtaining 55 distributions (Supplementary materials). Then, we selected the NTT SNPs, i.e. the Neanderthal-Tag SNPs in the Top 1% of each distribution (data file S4).

#### The biological implications of Neanderthal introgression

Given the list of genes overlapping the Neanderthal introgressed regions harbouring the NTT SNPs and the list of genes directly harbouring the NTT SNPs, we performed different enrichment tests with the online tool EnrichR (*60, 61*). Particularly, we searched for significant enrichments compared to the human genome using the EnrichR collection of database, e.g. dbGaP (*62, 63*), Panther 2016 (*64*), HPO (*65*) and KEGG 2016 (*66*–*68*) (data file S4). We then investigated known direct associations between the Neanderthal alleles of the NTT SNPs and phenotypes, by looking in the GWAS and PheWAS catalogues (*32, 33*) and by applying the PheGenI tool (*69*) (Supplementary Data 5). We used the circos representation as in Kanai et al. (*70*), to highlight different sets of NTT SNPs (Figure 4F).

## Acknowledgments

**General**: We would like to thank St Hugh’s College and the Department of Zoology for facilitating the visits of A.R. and S.A. to the University of Oxford and the PhD programs of the University of Pavia and University of Turin for supporting these visits; the High Performance Computing Facility of the Oxford University and CINECA for the computational resources, the programming assistance and advices given during this project; the SU.VI.MAX for the access to the FST estimates of their unpublished work (C.D., J.G., P.G.); Tony Capra for sharing the list of Neanderthal introgressed regions in humans; Ryan Daniels and Miguel Gonzales Santos for the computing advices during the early stages of this project; Luca Alessandri for his comments on the archaeological context of the Bronze Age in Italy and surrounding regions; Simonetta Guarrera for technical support (C.D.G., G.M.); the National Alpini Association (Associazione Nazionale Alpini) for their help in collecting Italian DNA samples at the 86^th^ national assembly in Piacenza in 2013, in particular Bruno Plucani, Giangaspare Basile, Claudio Ferrari and the municipality of Piacenza (A.O., A.A.). Finally, the authors would like to acknowledge all the people that donated their DNA and made this work possible.

## Author Contributions

A.O., A.A., G.M., F.M., and C.C. conceived the idea for the study; A.R., S.A., F.M. and C.C. performed, devised or supervised the analyses; A.A., A.O., F.B. and V.L.P. provided reagents for the genotyping of novel samples; all the authors contributed to this study providing data, computational facilities, or other resources; A.R., S.A., F.M. and C.C. wrote the manuscript with inputs from co-authors.

## Data and materials availability

Requests for accessing previously published data should be directed to the corresponding author of the publications where they were originally presented. Enquiries for unpublished data should be directed to Mait Metspalu (Genotype data provided by the Estonian Biocentre), Simon Myers (FST estimates among clusters in Spain), Christian Dina (FST estimates among clusters in France). Data genotyped as part of this project and presented here for the first time (135 Italian samples and 6 samples from Albania, genotyped on the Infinium Omni2.5-8 Illumina beadchip) can be downloaded at the following webpage: XXX

**Fig. S1.**
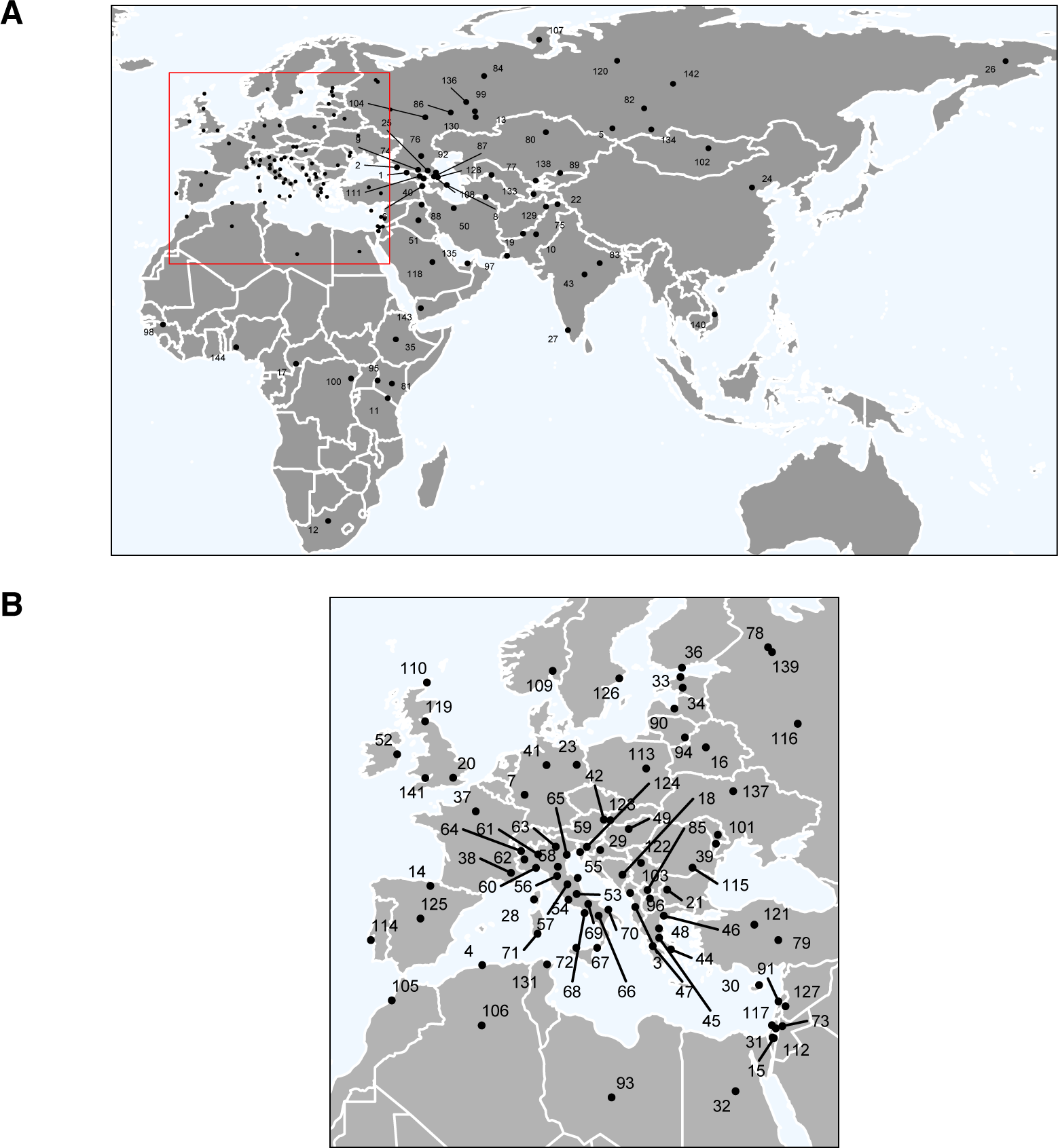
Geographic location of populations included in FMD and HDD. A) European, North African and Western Eurasia samples; B) World-wide samples. Numbers as in table S1.

**Fig. S2.**
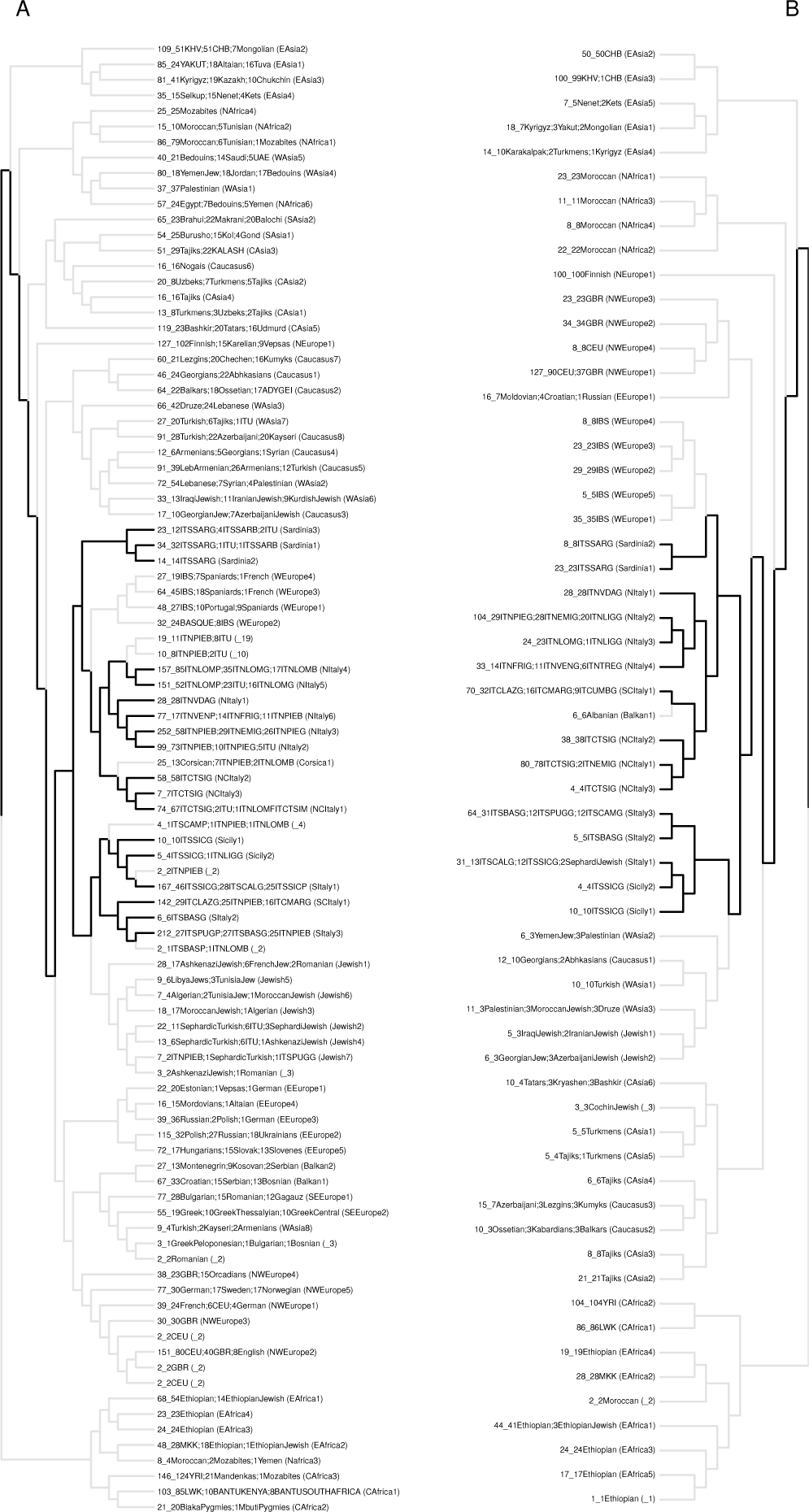
fineSTRUCTURE dendrogram of all the 4,852 (A, FMD) and 1,641 (B, HDD) samples. Each tip of the dendrograms represents a group of individuals with similar copying vectors. The first number of each tip label refers to the total number of individuals in the cluster. This value is followed by “_” and the name of the three most representative geographically-assigned populations, each with its number of samples. At the end, within brackets, the name given to the cluster. Thick lines in black refer to the Italian clusters. The details of cluster assignation are reported in data file S2.

**Fig. S3.**
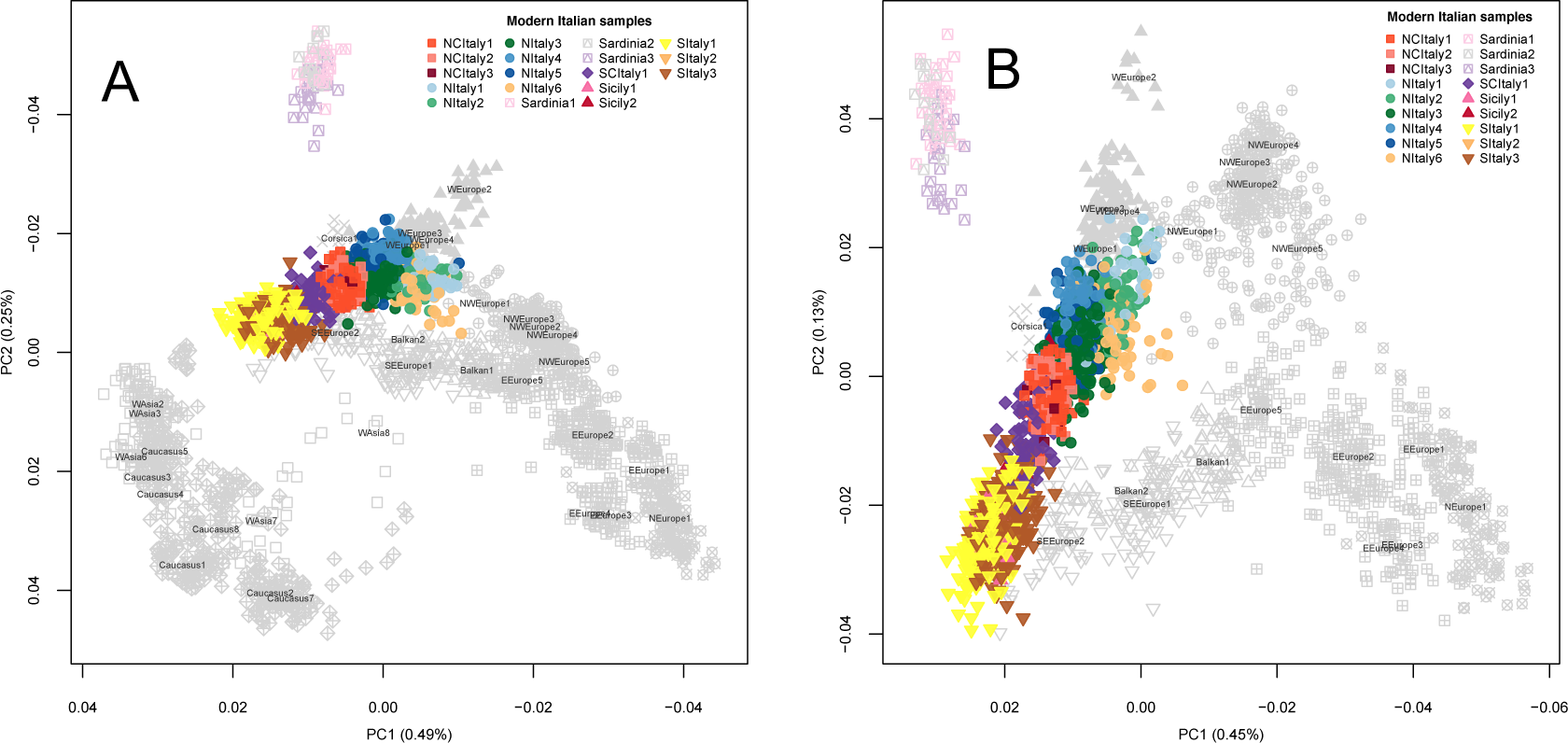
Allele frequency Principal Components Analysis (PCA) of modern samples (genotype based). A) PCA of 3,057 modern samples included in Eurasian CP/fS inferred clusters; all the samples are labelled and coloured as in Fig. 1A. B) PCA of 2,469 modern European samples as displayed from the dendrogram resulting from CP/fS (Fig. 1A).

**Fig. S4.**
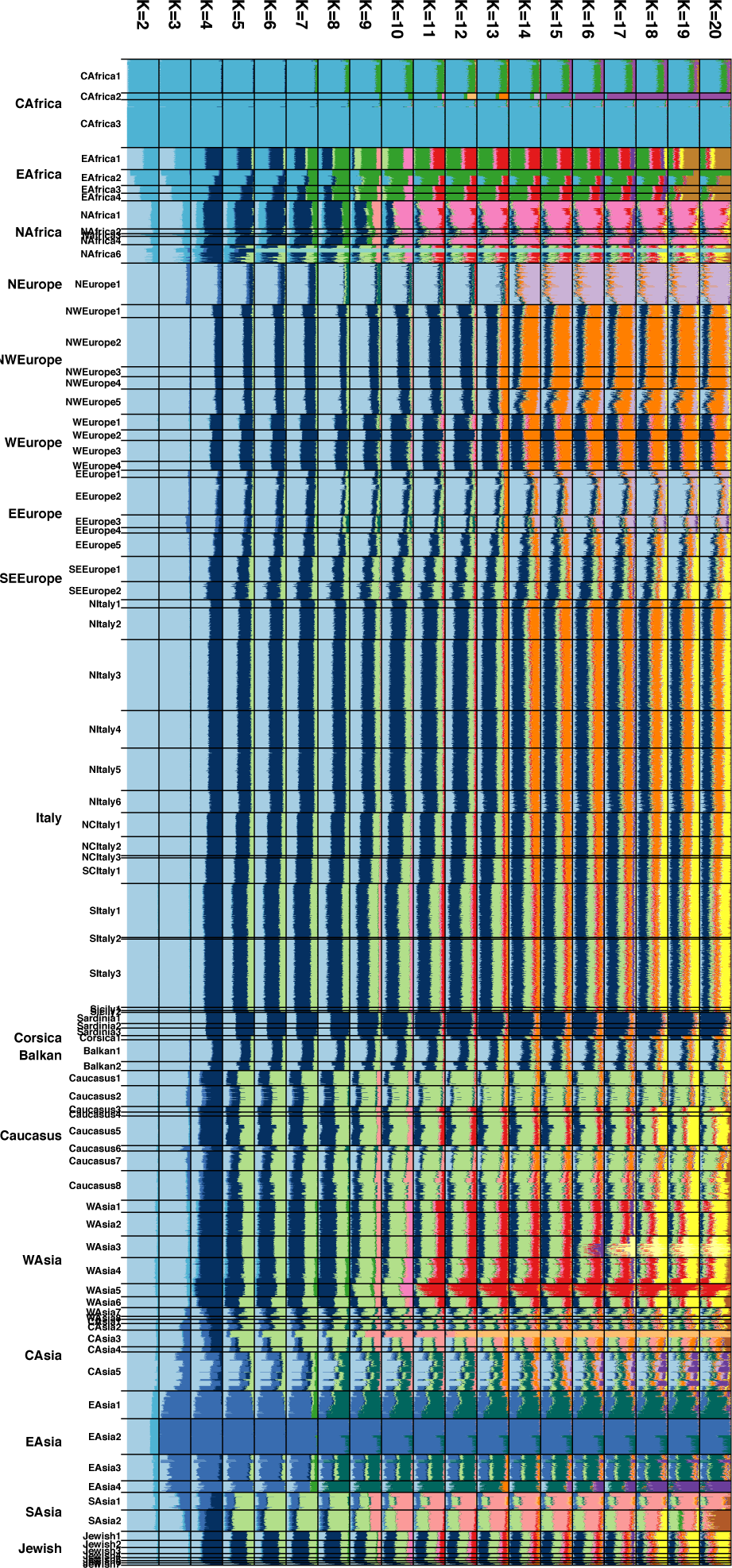
Individual-level ADMIXTURE analysis of modern samples. Samples are grouped according to the genetic clusters inferred by the CP/fS pipeline and named as in fig. S2.

**Fig. S5.**
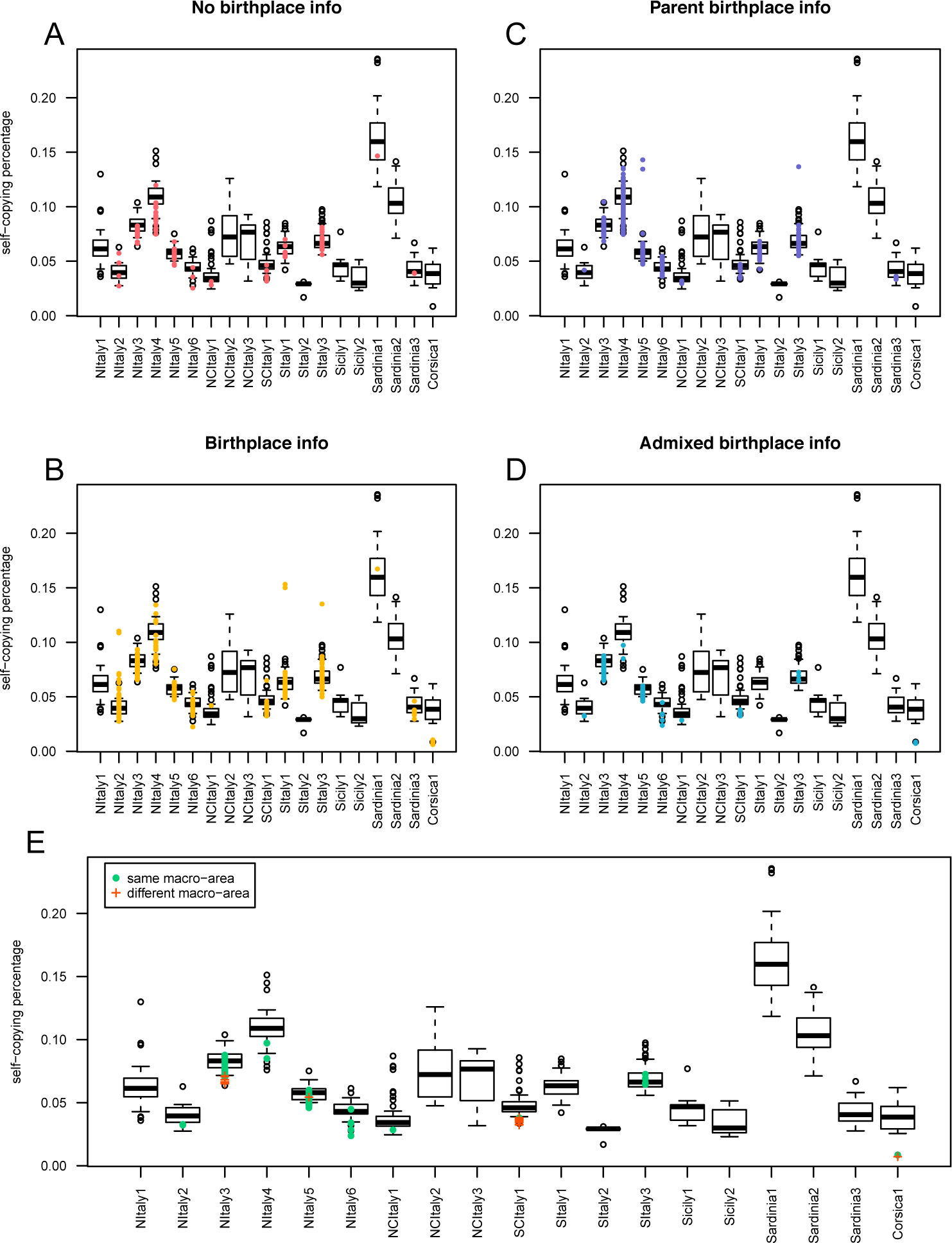
“Cluster self-copy” analysis. Box plots refer to the distributions of the self-copying vectors for each cluster for samples with same birthplace region for the four grandparents; coloured points refer to individual samples with other/no information; outliers are indicated as white circles. Coloured points refer to: A) subjects with no information available on their place of birth (red); B) subjects with only their own birthplace information (yellow); C) subjects with parents birthplace information (violet); D) subjects with “mixed” parental ancestry (parents from different regions) (blue); E) same as in D), red crosses identify individuals with parents born in different macro-areas (North and South Italy) indicated as suffix in each Italian population (table S1), while green dots refer to samples with parents born in the same macro-area.

**Fig. S6.**
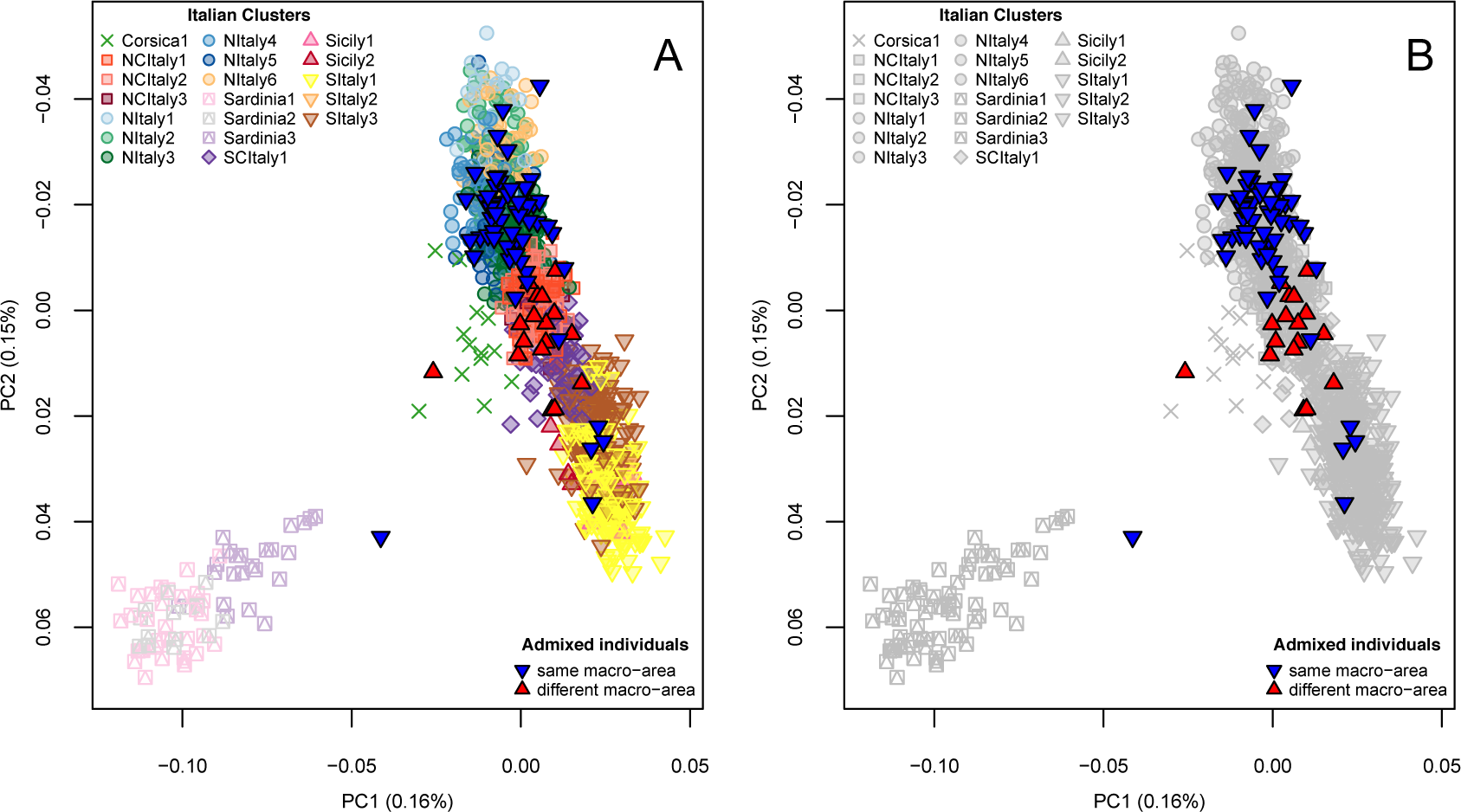
PCA with Admixed Italian individuals. Individuals with parents known to be born in two different macro-areas (see Materials and Methods, Supplementary materials - Cluster Self-Copy analysis) are plotted in red together with all the other Italian individuals, these coloured according either to the clusters they belong to (A) or in grey (B). Macro-areas are separated in Northern and Southern, where the central regions of Tuscany and Emilia are considered as part of the Northern macroarea and Latium, Abruzzo, Marche and Sardinia were considered as part of the Southern macro-area.

**Fig. S7.**
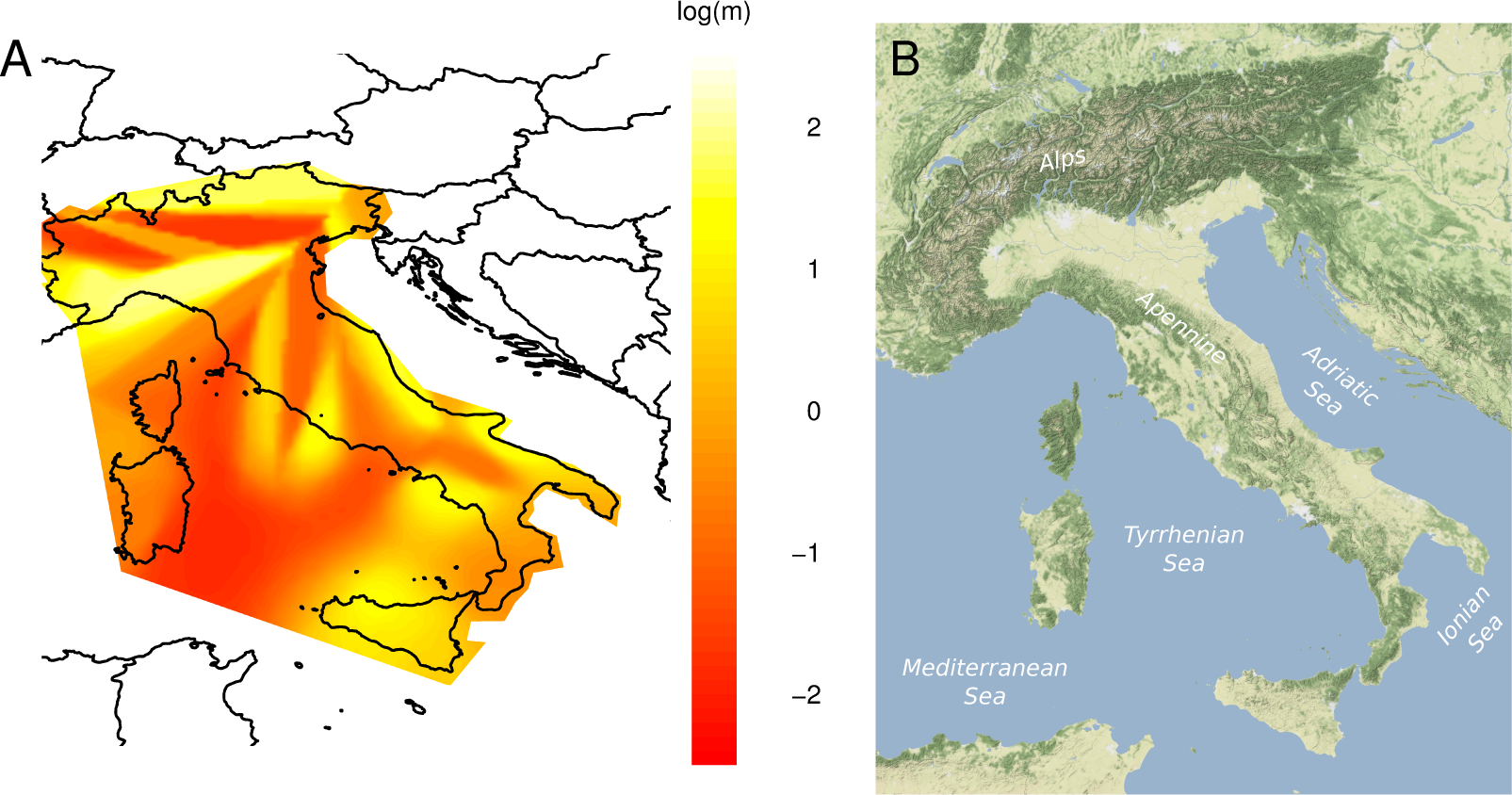
Results of the EEMS analysis on Italy-only populations. A) Colours represent the log10 scale of the effective migration rate from low (red) to high (yellow). Samples as reported in table S1. B) Physical map of Italy.

**Fig. S8.**
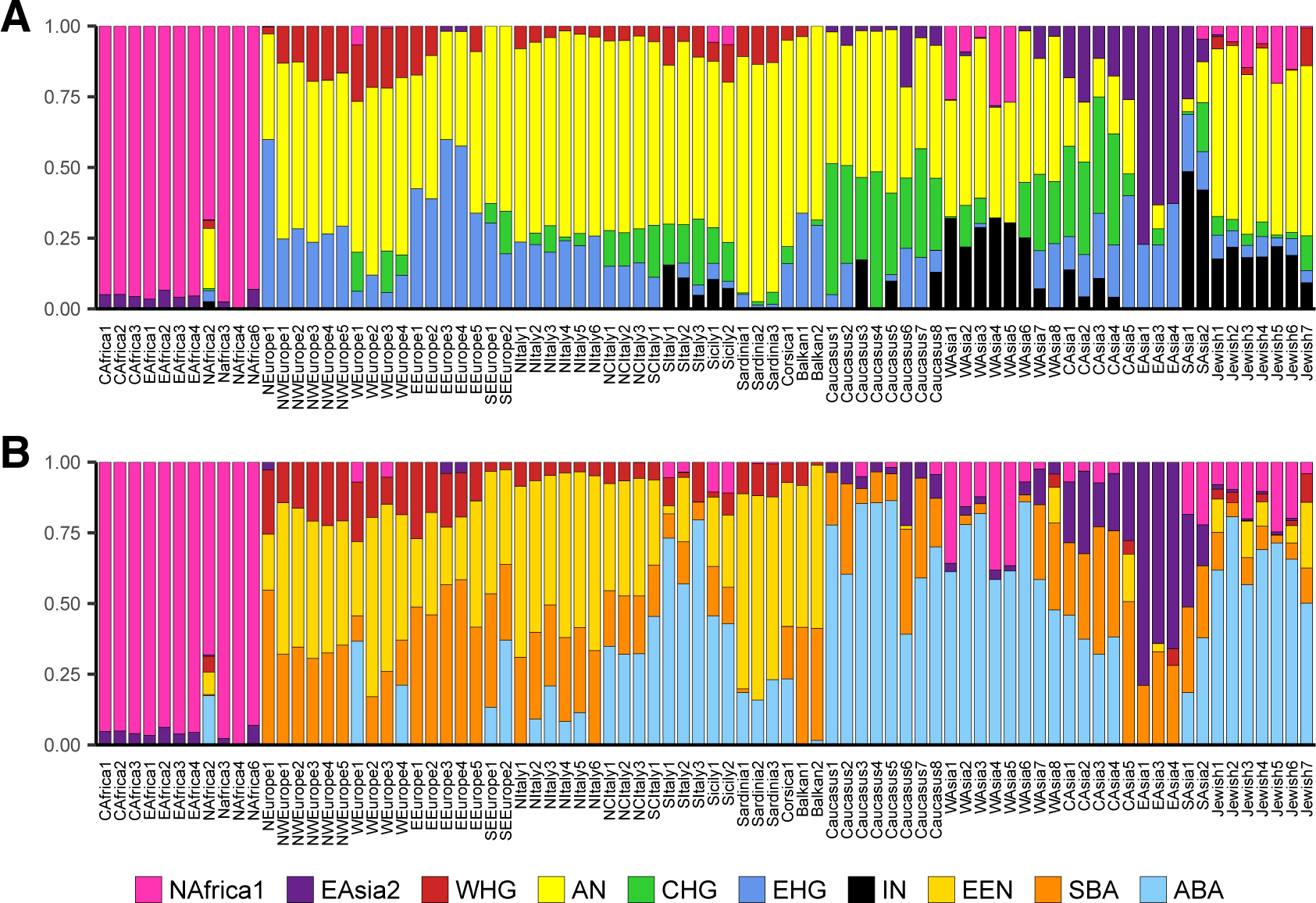
CP/NNLS results for *Ultimate* and emphProximate sources for all modern clusters. A) *Ultimate* (A) and *Proximate* (B) sources analysis reporting all modern Eurasian and African clusters and including WHG among the sources (main text; Supplementary Material).

**Fig. S9.**
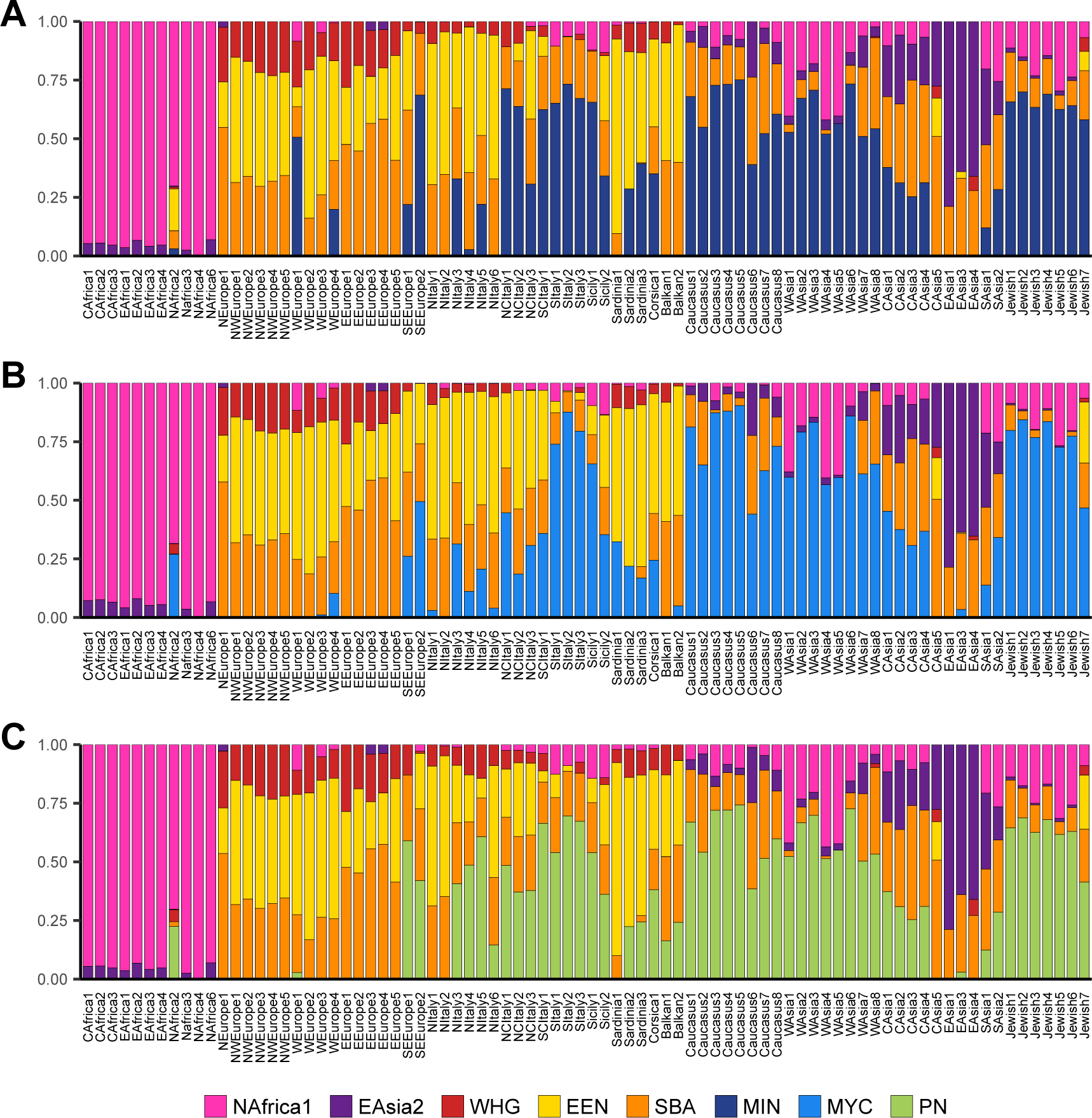
CP/NNLS results for *emphProximate* sources for all modern clusters using alternative SEE sources. *Proximate* sources analysis replacing ABA with alternative SEE sources: A) Minoan, MIN: B) Myce-naean, MYC: C) Peloponnese Neolithic, PN. In all the analyses, WHG was included among the possible sources (Supplementary Material).

**Fig. S10.**
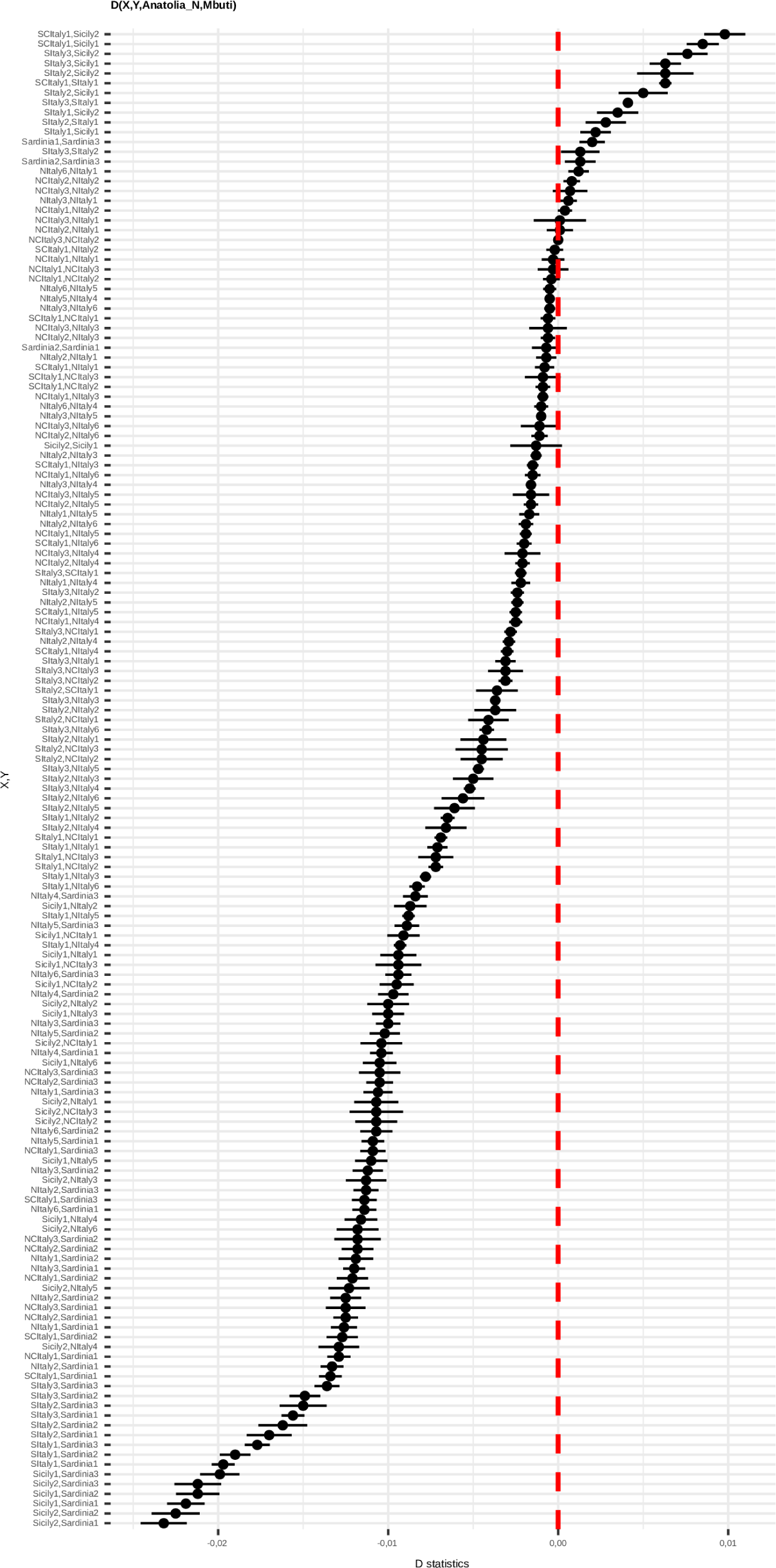
D statistics in the form D(X,Y, AN,Mbuti) for all the possible pairs of Italian clusters.

**Fig. S11.**
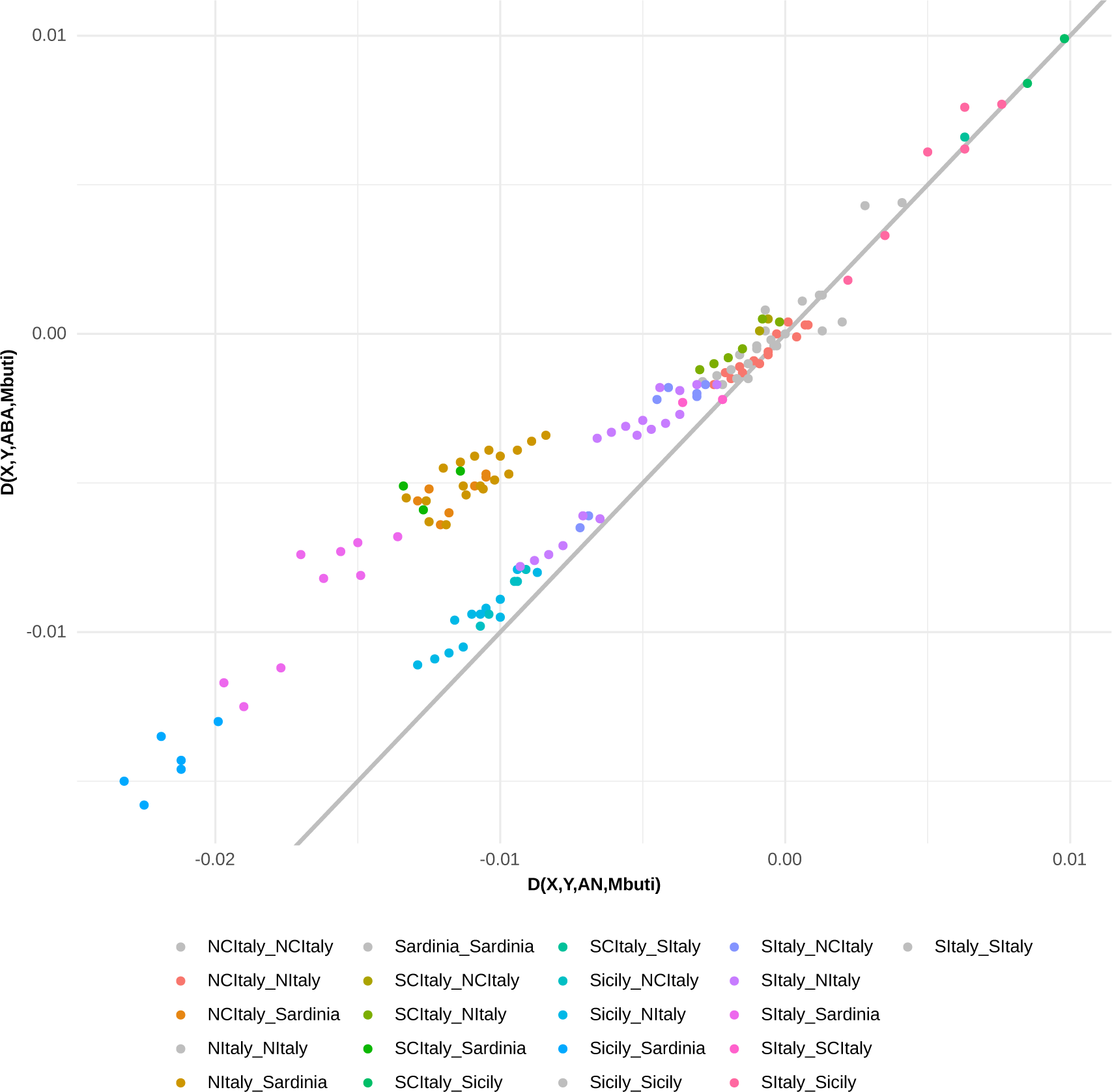
Comparison of AN and ABA affinity to Italian clusters using D-statistics. Scatter plot of D(Ita1, Ita2, AN, Mbuti) and D(Ita1,Ita2,ABA,Mbuti) for all the Italian clusters. Points for pairs of clusters from the same (grey points) or closely related geographic location fall in proximity of the grey line, reflecting a similar affinity to AN (x-axis) and ABA (y-axis). Comparisons of clusters from NItaly/Sardinia and SItaly/Sicily fall above the grey line, reflecting a closer affinity of the latter to ABA.

**Fig. S12.**
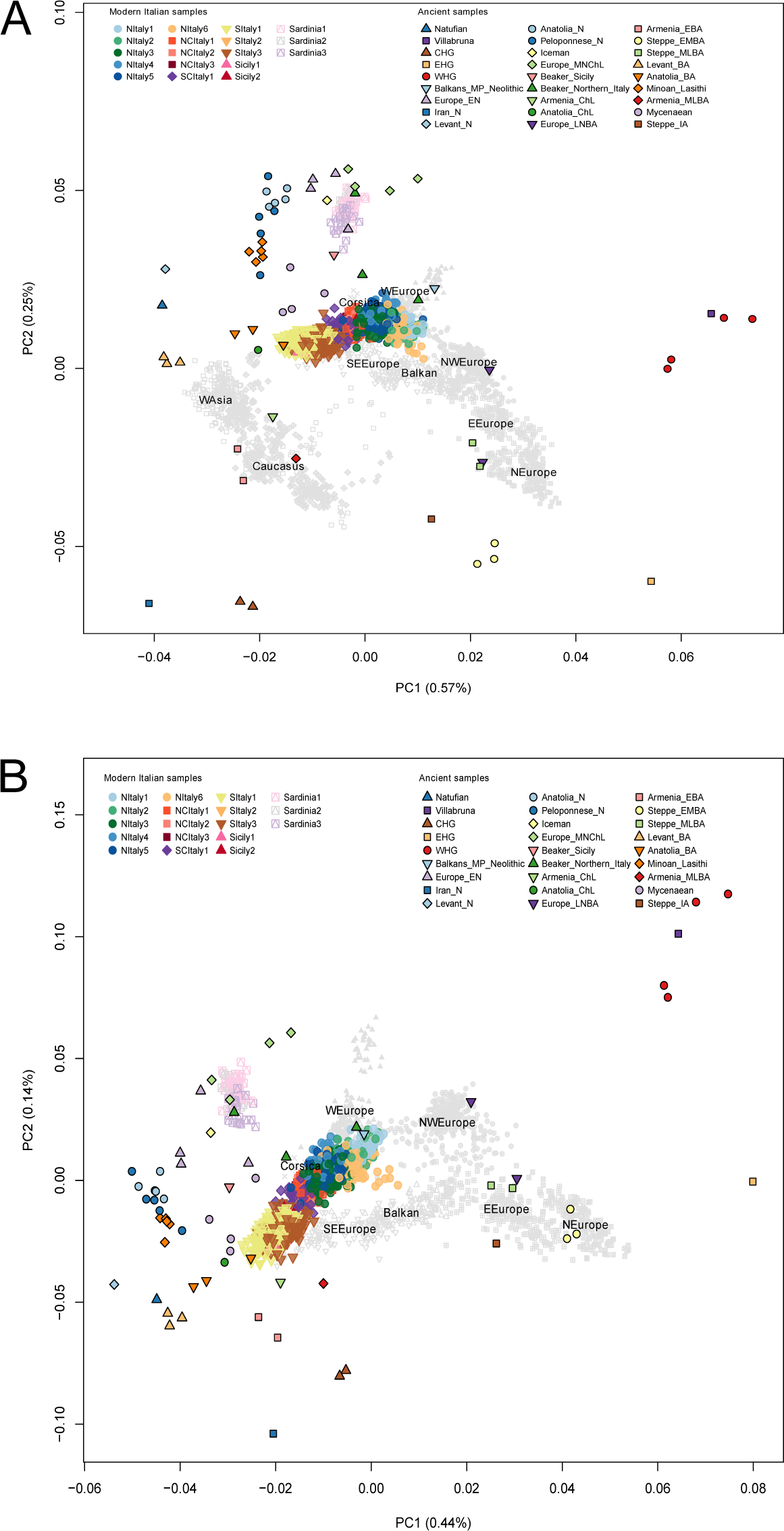
Principal component analysis projecting 63 ancient individuals onto the components inferred from modern individuals. A) Principal component analysis projecting 63 ancient individuals onto the components inferred from 3,282 modern individuals assigned, through a CP/fS analysis, to European West Asian and Caucasian clusters (data file S2). B) Principal component analysis projecting 63 ancient individuals onto the components inferred from 2,469 modern individuals assigned, through a CP/fS analysis, to European clusters (data file S2). The labels are placed at the centroid of the macroarea. The centroids are calculated by computing the means of the coordinates of individuals in modern clusters within each macroarea.

**Fig. S13.**
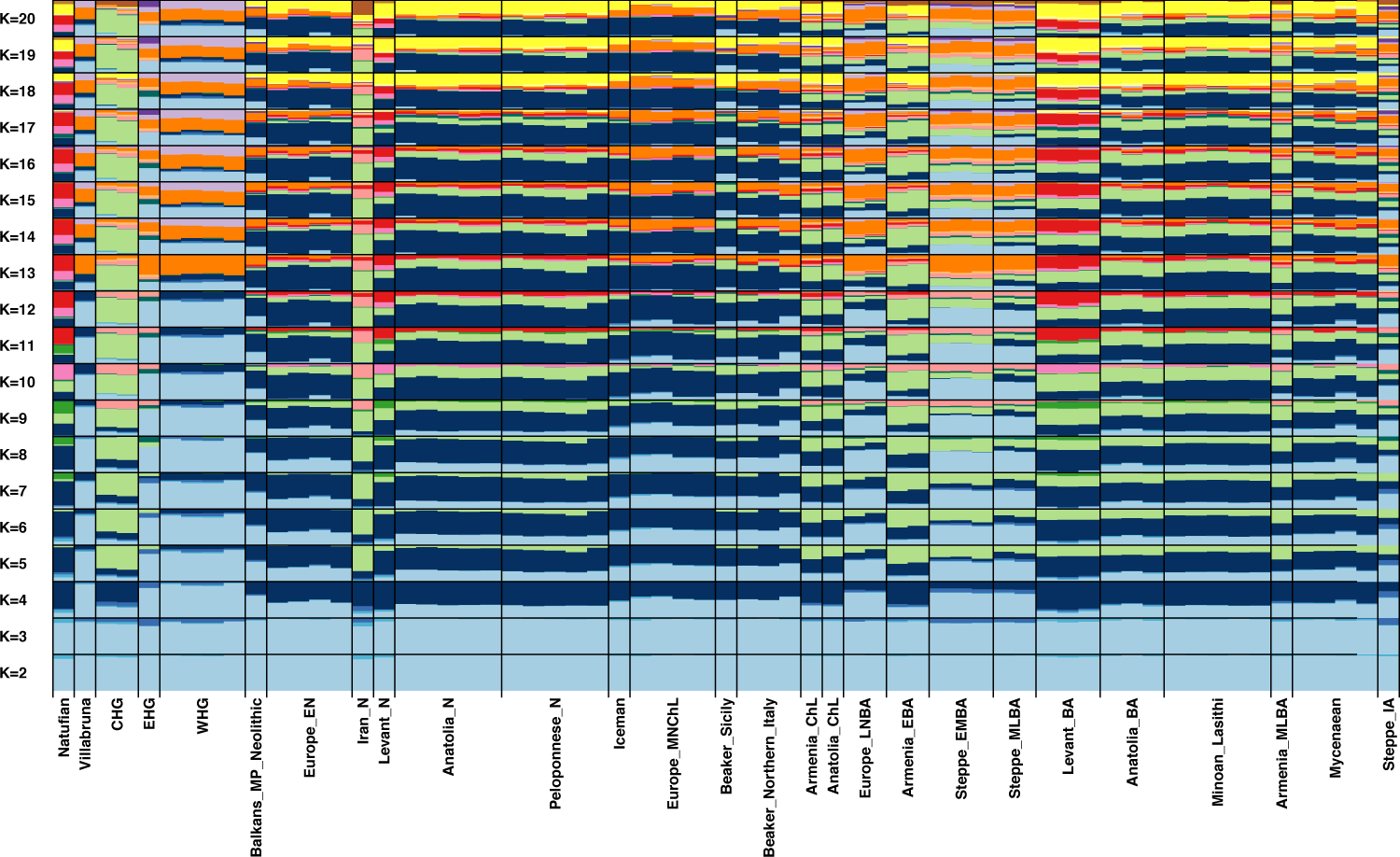
ADMIXTURE analysis of 63 ancient samples. Ancestral allele frequencies were inferred from ten different ADMIXTURE runs on 4,606 modern samples and projected onto the ancient samples. Each bar represents an individual grouped into ancient groups (data file S1).

**Fig. S14.**
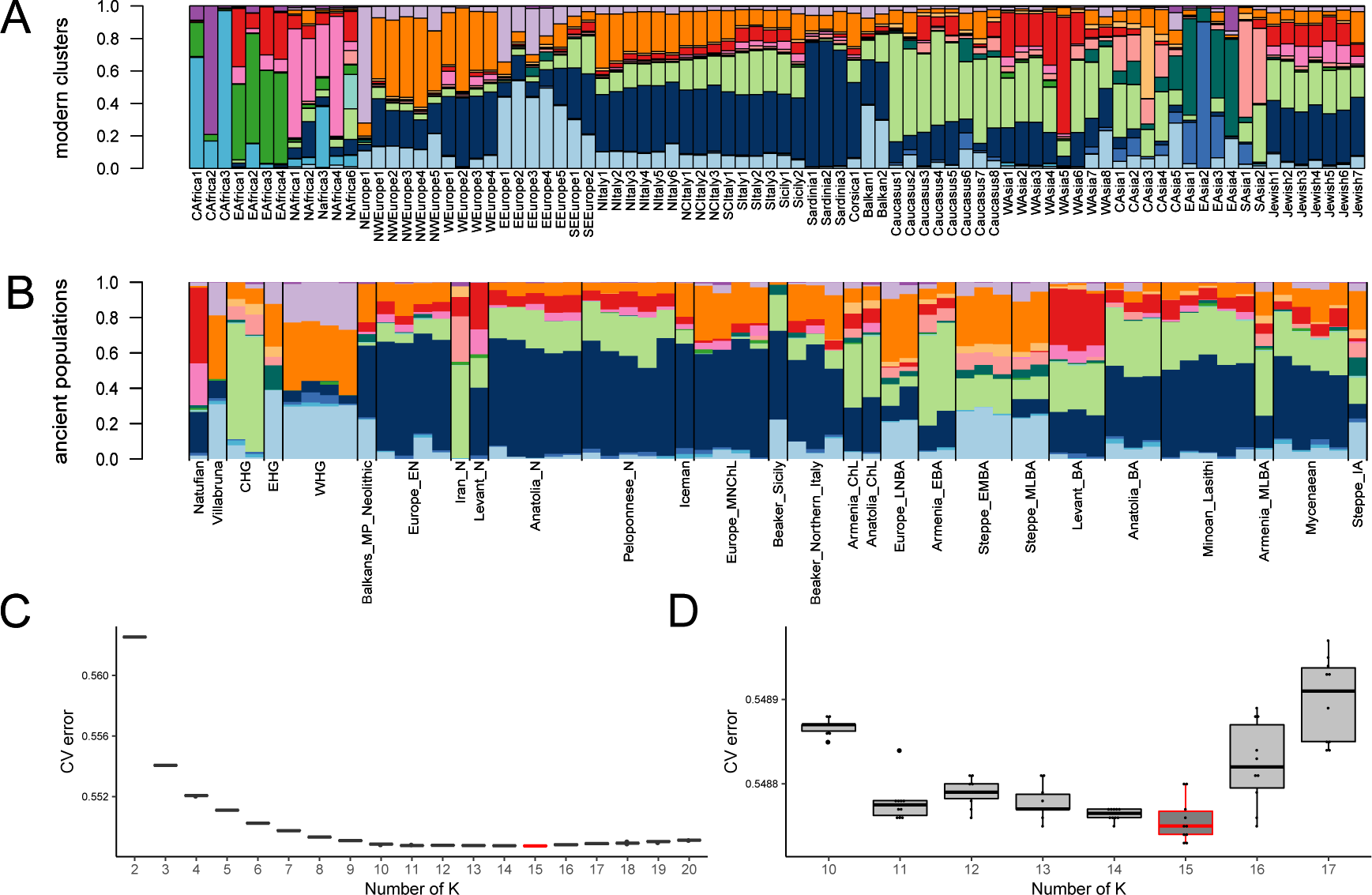
ADMIXTURE analysis of 63 ancient samples and 4,606 modern samples for K=15. A-B) Results of the ADMIXTURE analysis as in fig. S4 and fig. S13 for K=15 including both modern (A) and ancient samples (B). C) Box plots of the ten CV-errors of each K from 2 to 20. D) Detailed box plots for the ten CV-errors for each K from 10 to 17.

**Fig. S15.**
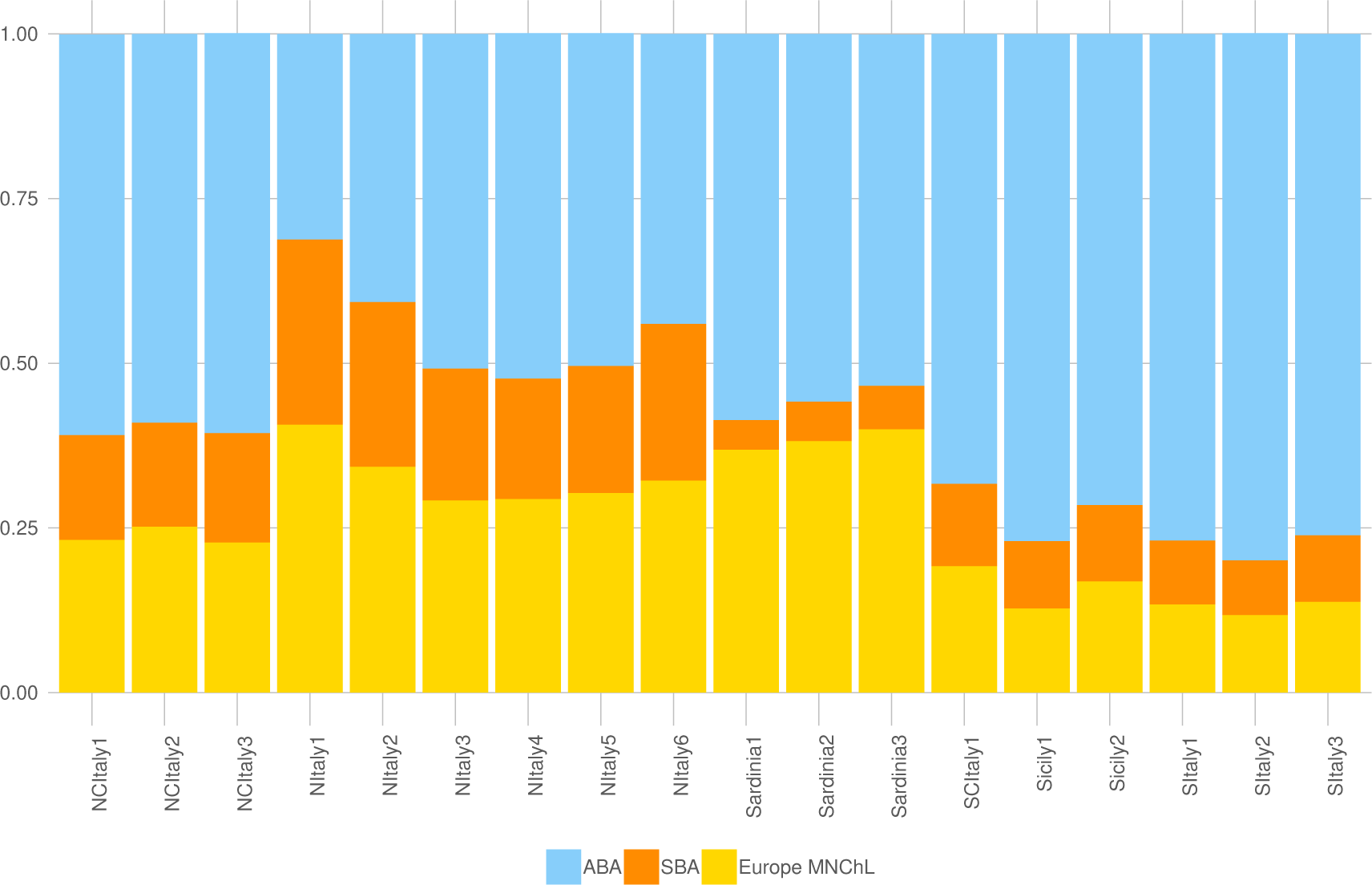
Mixture proportions on modern Italian clusters inferred by qpAdm as a combination of ABA, SBA and European Middle-Neolithic/Chalcolithic. For each tested cluster, we have evaluated all the possible combinations of N “left” sources with N={2‥5}, and one set of right/left Outgroups (Supplementary materials).

**Fig. S16.**
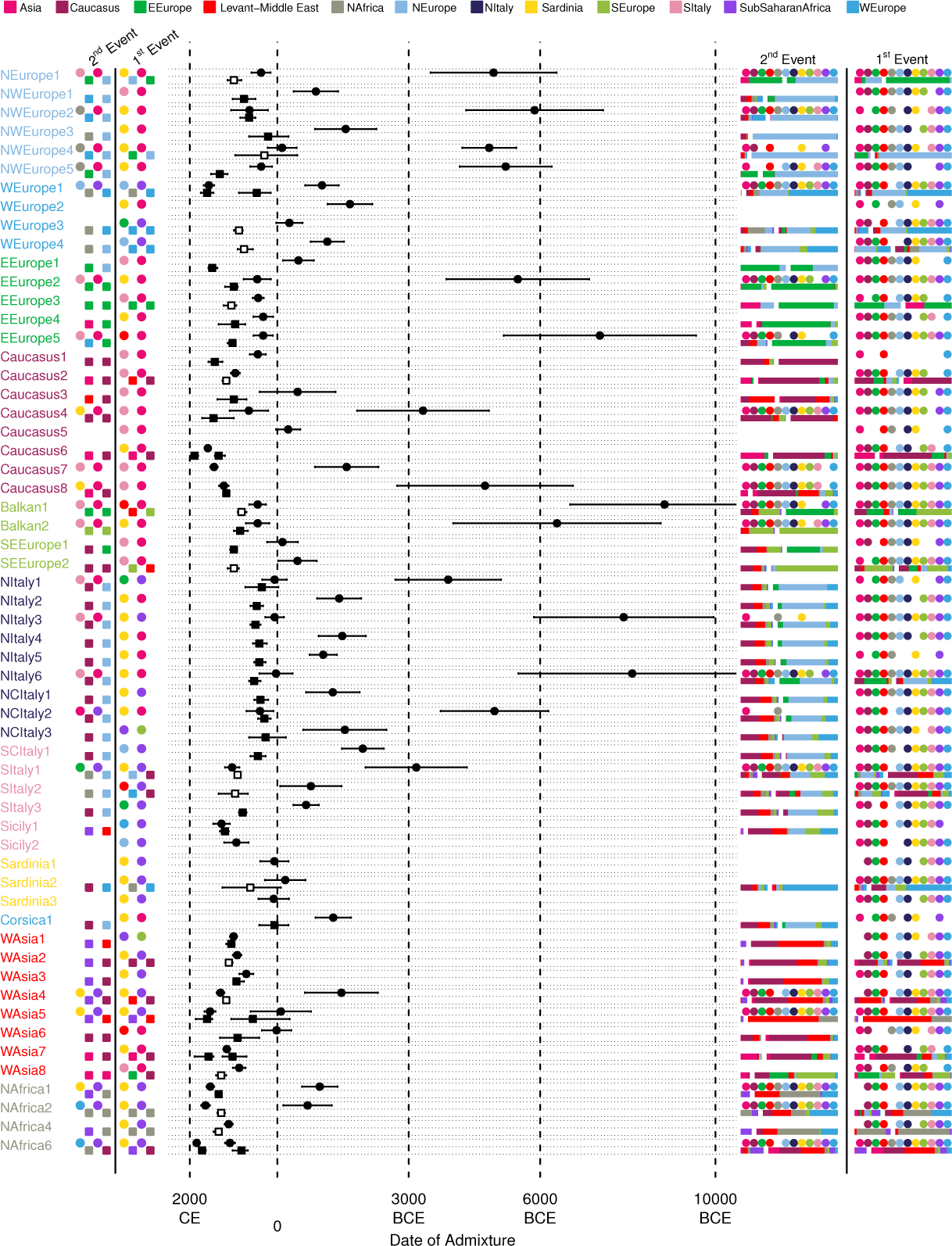
GT and MALDER analyses for all the Eurasian and North African clusters. Dates of the events inferred by “noItaly” GT (squares) and MALDER (circles) for clusters as in Fig. 1A and data file S2 are reported in the central part of the plot; lines encompassed the 95% CI for GT and ±1 Standard Error for MALDER. GT events were distinguished in “one date” (black squares; 1D in table S7), “one date multiway” (white squares; 1MW) or “two events” (two black squares; 2D). The best sources are indicated in a staggered way as circles and squares for MALDER and GT, respectively (“1 ^st^ /2 ^nd^ event” columns, on the left; four sources are highlighted for 1MW events). Colours refer to the ancestry to which the sources were assigned (see Materials and Methods; Supplementary materials). We additionally included a sub-Saharan African ancestry comprising CAfrica and EAfrica clusters (Fig. S2, data file S2). GT sources for single date events are plotted in the column “2 ^nd^ event”, as overlapping with second events detected by MALDER. The composition of the sources for GT and the geographical regions of the sources in MALDER, for which no significant differences in the amplitude of the fitted curve were found, are reported in the “1 ^st^ /2 ^nd^ event” columns on the right. GT sources are divided by a white space; the length of the bars indicates the contribution of each source; for 1MW events, two bar plots are indicated in the “1 ^st^ /2 ^nd^ event” columns on the right.

**Fig. S17.**
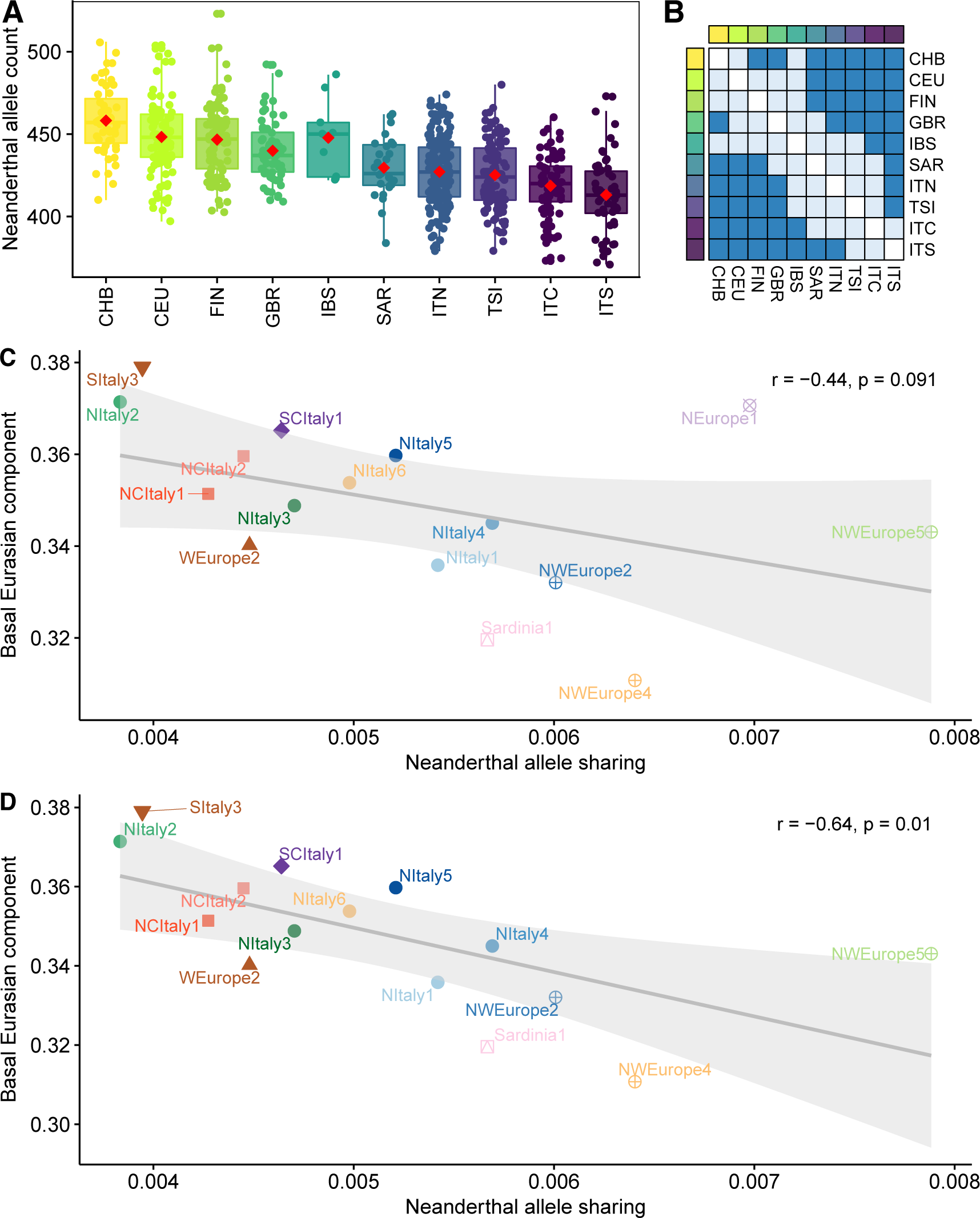
Exploring the relationship between Neanderthal ancestry and admixture with African sources. Same as in Fig. 4A, B, C but removing either the individuals belonging to clusters where the GT analysis identified signatures of African admixture (clusters SItaly1, SItaly2, Sicily1, Sardinia2, NWEurope3, WEurope1, WEurope3 and WEurope4, Figure 3 and fig. S16) or the whole set of the clusters listed above (see Supplementary materials). Specifically: A) Neanderthal allele counts in individuals from Eurasian populations, on 3,969 LD-pruned Neanderthal tag-SNPs; B) Matrix of significances based on Wilcoxon rank sum test between pairs of populations including (lower triangular matrix) and removing (upper) outliers (dark blue: adj p-value < 0.05; light blue: adj p-value > 0.05). C) Correlation between Neanderthal ancestry proportions and the amount of Basal Eurasian ancestry in European clusters. D) Same as C) but removing the cluster NEurope1 (see Supplementary Materials). Clusters with less than 10 individuals were excluded in C and D.

**Fig. S18.**
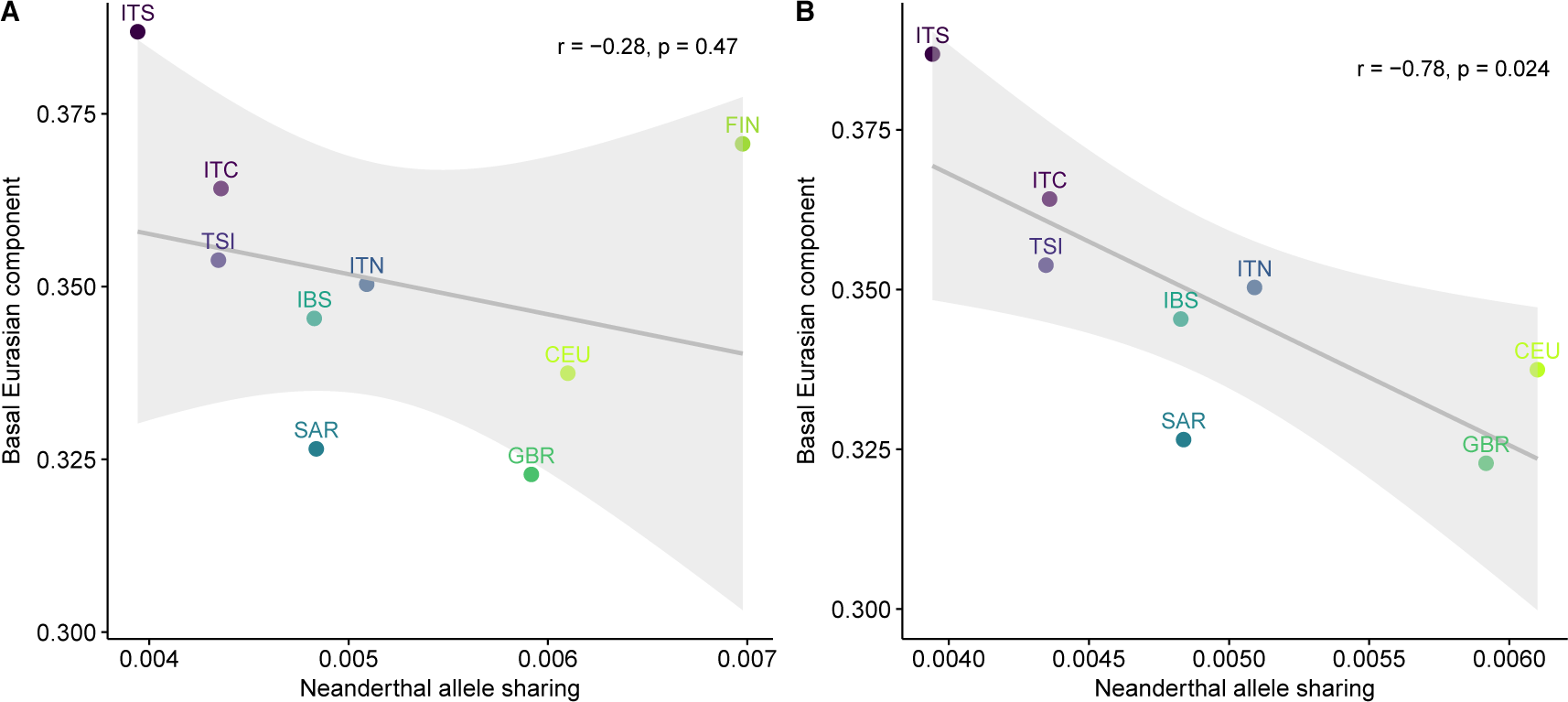
Correlation between the proportion of Neanderthal allele sharing and the amount of ancestry derived from a Basal Eurasian population in European populations. A) Correlation considering FIN (Finnish in Finland) population. B) Correlation excluding FIN (Finnish in Finland) population (see Materials and Methods, Supplementary materials).

**Fig. S19.**
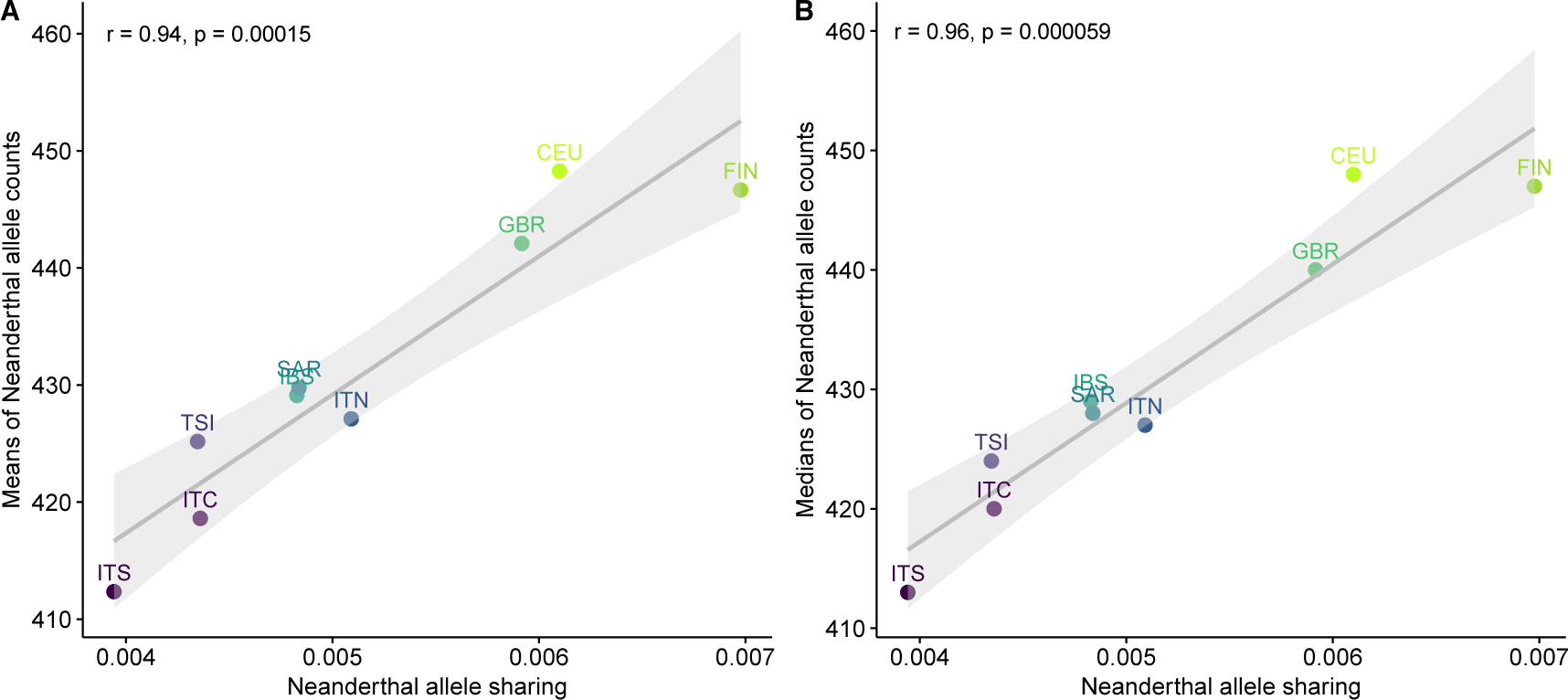
Correlation between the proportions of Neanderthal allele sharing computed with F4-ratio and the counts per population of Neanderthal alleles in European populations. A) Correlation between the proportions of Neanderthal allele sharing computed with F4-ratio and the means per population of Neanderthal allele counts. B) Correlation between the proportion of Neanderthal allele sharing computed with F4-ratio and the medians per population of Neanderthal allele counts.

**Fig. S20.**
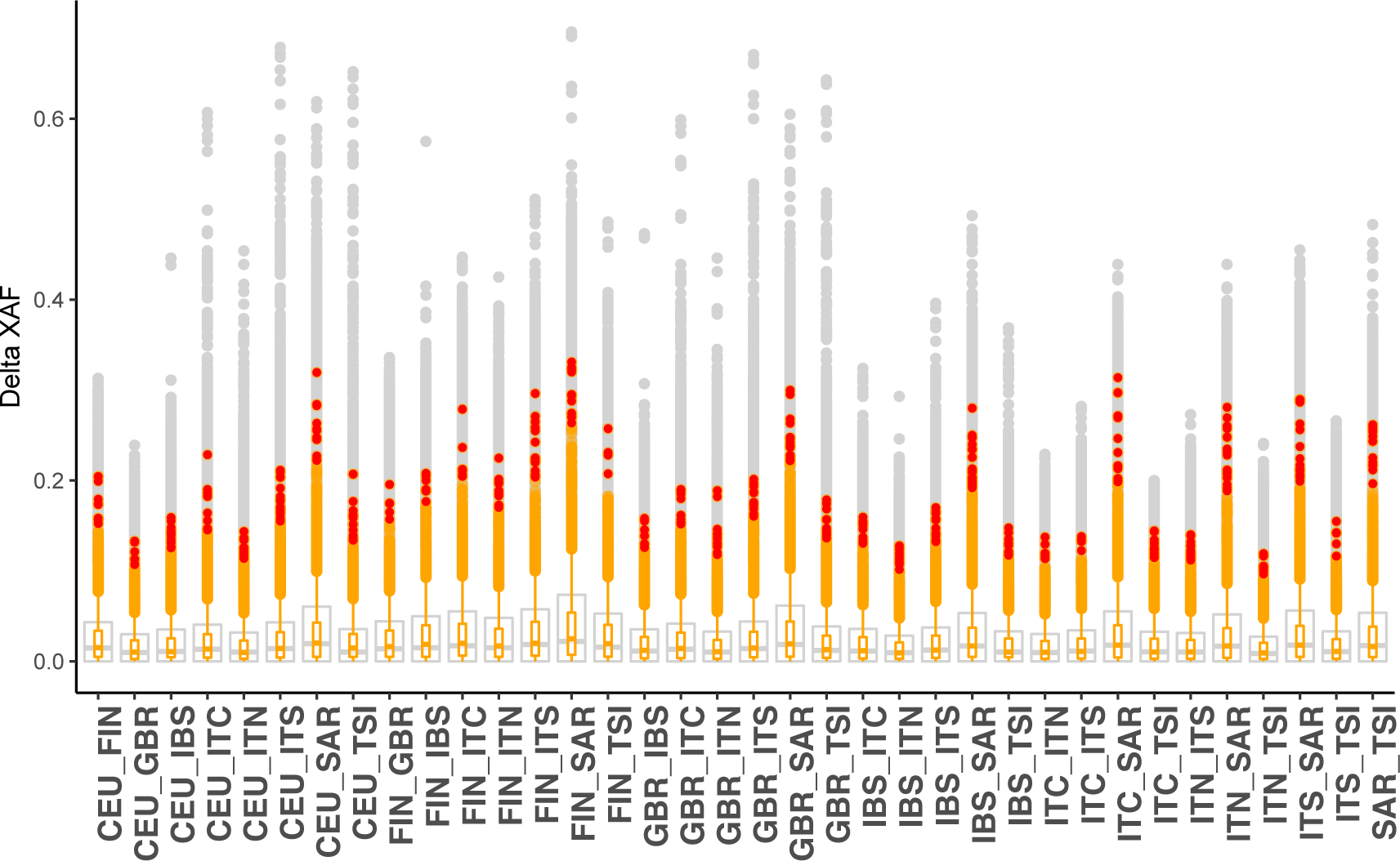
Absolute allele frequency differences (ΔXAF, where X is the minor allele for each SNP or the Neanderthal allele when considering Neanderthal regions tag-SNPs) for each pair of European populations. We reported in grey the boxplot representing the total distributions of the variants, and in orange the distribution of Neanderthal inherited variants. The red dots are the Neanderthal SNPs in the top 1% of the distributions, as also reported in data file S4.

